# *Bacopa monnieri* phytochemicals as promising BACE1 inhibitors for Alzheimer’s Disease Therapy

**DOI:** 10.1101/2024.09.19.613992

**Authors:** Satyam Sangeet, Arshad Khan

## Abstract

Alzheimer’s disease (AD) remains a formidable challenge, necessitating the discovery of effective therapeutic agents targeting β-site amyloid precursor protein cleaving enzyme 1 (BACE1). This study investigates the inhibitory potential of phytochemicals derived from *Bacopa monnieri*, a plant renowned for its cognitive-enhancing properties, in comparison to established synthetic inhibitors such as Atabecestat, Lanabecestat, and Verubecestat. Utilizing molecular docking and advanced computational simulations, we demonstrate that Bacopaside I exhibits superior binding affinity and a unique interaction profile with BACE1, suggesting a more nuanced inhibitory mechanism. Our findings highlight the promising role of *Bacopa monnieri* phytochemicals as viable alternatives to synthetic drugs, emphasizing their potential to overcome limitations faced in clinical settings. Furthermore, the development of the SIMANA (https://simana.streamlit.app/) platform enhances the visualization and analysis of protein-ligand interactions, facilitating a deeper understanding of the dynamics involved. This research not only underscores the therapeutic promise of natural compounds in AD treatment but also advocates for a paradigm shift towards integrating traditional medicinal knowledge into contemporary drug discovery efforts.

## Introduction

Alzheimer’s disease (AD) is a neurodegenerative disorder that ranks among the most prevalent causes of dementia globally, primarily affecting the aging population^1,2^. AD is characterized by the progressive deterioration of cognitive functions, memory loss, and the gradual impairment of daily activities. At the molecular level, the disease’s hallmark features include the accumulation of extracellular amyloid plaques and intracellular neurofibrillary tangles in regions of the brain such as the hippocampus and cortical gray matter^3,4,5^. These plaques are primarily composed of amyloid-beta (Aβ) peptides, which arise from the sequential cleavage of the amyloid precursor protein (APP) by two key enzymes—β-secretase (BACE1) and γ-secretase. The β-secretase enzyme, in particular, plays a critical role in the initial and rate-limiting step of this proteolytic process, making it a prime therapeutic target for AD.

BACE1 (β-site of Amyloid Precursor Protein Cleaving Enzyme) is a membrane-bound aspartyl protease that plays a critical role in the production of amyloid-beta (Aβ) peptides, central to Alzheimer’s disease pathology^6,7,8,9^. Its active site contains two essential aspartate residues, Asp32 and Asp228 (Fig. 1), which coordinate a water molecule, facilitating the nucleophilic attack necessary for peptide bond hydrolysis. A unique feature of BACE1 is the flap region (residues 67–77), a hairpin loop that partially covers the active site cleft. This flap remains open in the enzyme’s inactive state^8,9^ but closes over the substrate or an inhibitor during catalysis, stabilizing the enzyme-substrate complex. Additionally, the 10s loop (residues 9–14) (Fig. 1), undergoes conformational changes that regulate substrate access to the active site. These structural features, particularly the dynamic nature of the flap and 10s loop, are key to understanding BACE1’s substrate specificity and catalytic mechanism, making them critical targets in the design of Alzheimer’s therapeutics.

**Figure 1:**
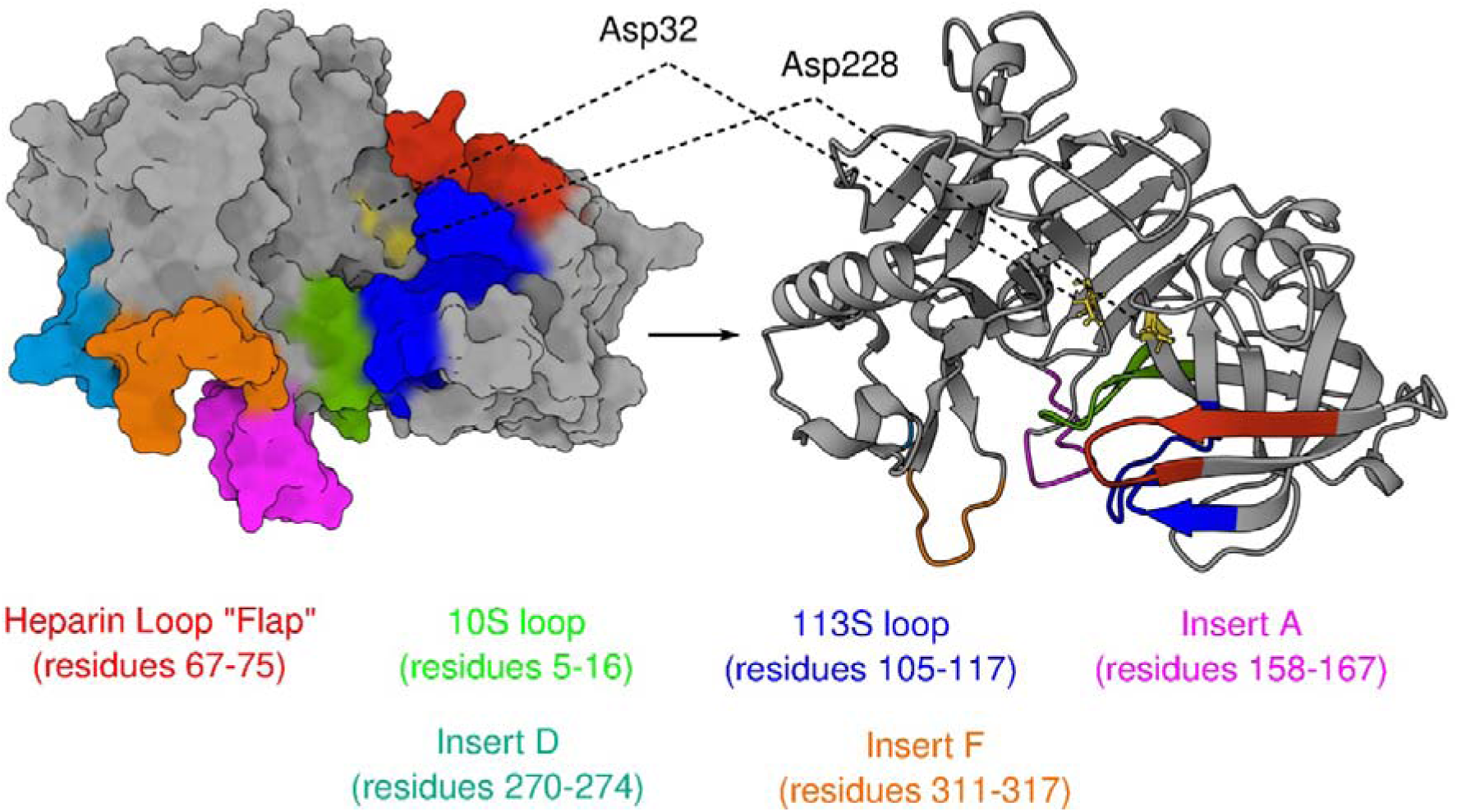
Structure of BACE1 in surface view (left) and cartoon view (right). Important regions of BACE1 are highlighted in their respective colour and residue range.

Molecular dynamics simulations approaches have elucidated how structural variations in regions such as inserts A, D, and F, along with the 10s loop, affect substrate binding and catalytic activity^10^. Various inhibitors have been developed to target BACE1, but their success has been limited by challenges such as off-target effects^11,12^. Atabecestat^13^, Lanabecestat^14,15^, and Verubecestat^16^ were among the most promising BACE1 inhibitors developed to target Alzheimer’s disease by reducing amyloid-beta (Aβ) production. These inhibitors progressed through clinical trials with hopes of curbing Aβ accumulation and slowing disease progression. However, despite their ability to effectively reduce Aβ levels, all three compounds failed in advanced clinical trials. Atabecestat was discontinued due to liver toxicity^17^, while Lanabecestat and Verubecestat were halted after showing no cognitive benefit in patients^18,19^. In particular, Verubecestat, which reached Phase III trials, revealed that although Aβ reduction occurred, the cognitive decline continued unabated, raising concerns about the complexity of Aβ’s role in Alzheimer’s pathology^20^. These failures highlight the challenges in translating BACE1 inhibition into an effective therapeutic strategy, as simply reducing Aβ levels may not be sufficient to alter the course of Alzheimer’s disease.

*Bacopa monnieri*, a traditional medicinal herb, has garnered significant attention for its neuroprotective properties, particularly in combating neurodegenerative disorders like Alzheimer’s disease^21,22,23,24^. Rich in bioactive compounds such as bacosides, flavonoids, and alkaloids, *B. monnieri’s* phytochemicals have demonstrated potent antioxidant, anti-inflammatory, and neuroprotective effects. It has been shown that *Bacopa monnieri* rejuvenates nerve cells and improves cognition and memory^25^. Studies have shown that *Bacopa monnieri* not only reduces Aβ deposition but also enhances cognitive function and neuronal health, making it a compelling natural candidate for Alzheimer’s therapy^26,27^. Moreover, we have shown that the phytochemicals of *Bacopa monnieri* has the potential to act as potent anti-neurodegenerative molecule by inhibiting the neurotrophins related neurological disorders^28^.In the current study, we have harnessed the therapeutic potential of *B. monnieri’s* phytochemicals to explore their efficacy in targeting Alzheimer’s-related neurodegeneration. By leveraging these compounds, we aim to investigate their role as promising alternatives or adjuncts to synthetic BACE1 inhibitors, potentially offering a safer and more holistic approach to mitigating cognitive decline and Aβ toxicity.

## Results and Discussion

### Phytochemical and Drug Selection

In this investigation, we selected a suite of eight biologically active phytochemicals, based on their reported efficacy in previous studies^29^. The molecular structures of these compounds are depicted in Fig. 2. To ensure accuracy, we cross-referenced the structural configurations and molecular masses of these phytochemicals against the PubChem database (https://pubchem.ncbi.nlm.nih.gov/)^30^. To benchmark the efficacy of the selected phytochemicals, we considered three known BACE1 inhibitors: Atabecestat, Lanabecestat and Verubecestat (Fig. 2).

**Figure 2:**
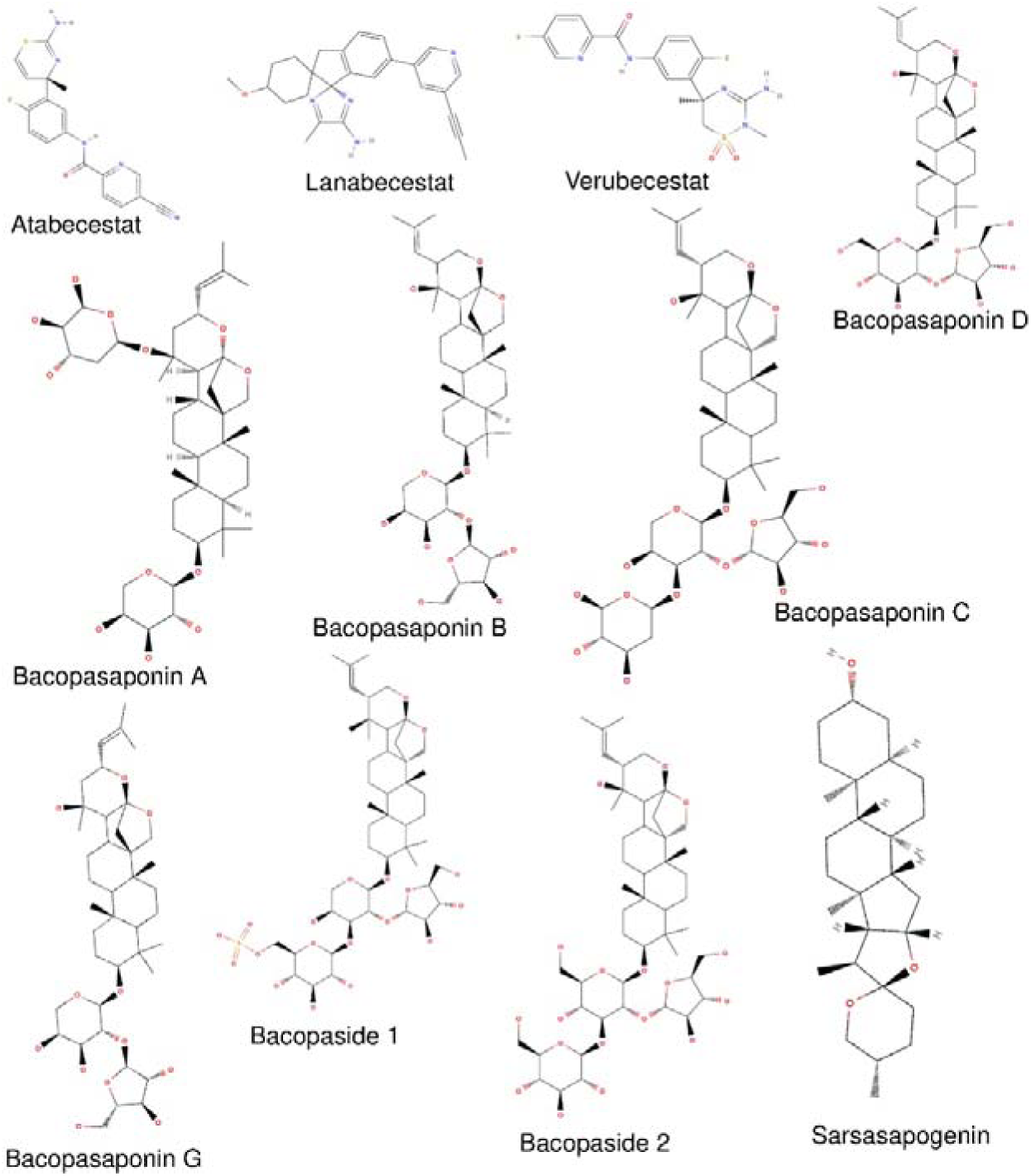
Known inhibitor of BACE1 along with the selected phytochemicals of *Bacopa monnieri*

### Molecular Docking

The molecular docking analysis comparing three established BACE1 inhibitors (Atabecestat, Lanabecestat, and Verubecestat) and the phytochemicals from *Bacopa monnieri* revealed intricate details about the ligand interaction. Atabecestat exhibited a binding energy of -7.5 kcal/mol (Table 1) and formed three hydrogen bonds with key residues: Asp228, Gly230, and Thr231 (Fig. 3a). Hydrophobic interactions with ten residues, including Ser35, Gly34, and Tyr71 (Table 2), contributed to the stability of the ligand-protein complex, suggesting a strong inhibitory potential. Lanabecestat showed the most favourable binding energy of -8.8 kcal/mol (Table 1), despite forming only one hydrogen bond with Thr72 (Fig. 3b). Its extensive hydrophobic interactions with residues like Asp32, Asp228, Gly34, and Tyr71 (Table 2) likely compensated for the limited hydrogen bonding, supporting its high binding affinity. Verubecestat, with a binding energy of -7.4 kcal/mol (Table 1), formed two hydrogen bonds with Asp228 and Thr231, directly interfering with the enzyme’s active site (Fig. 3c). It also exhibited hydrophobic interactions with nine residues, including Gly34 and Tyr71 (Table 2), further stabilizing the ligand-enzyme complex. Among the phytochemicals, Bacopasaponin A displayed a binding energy of -8.6 kcal/mol (Table 1), comparable to Lanabecestat. It formed an extensive network of five hydrogen bonds, including with Asp228, Gly34, and Thr231 (Fig. 3d), and interacted hydrophobically with eleven residues, such as Tyr71, Leu30, and Phe108 (Table 2). This combination of hydrogen bonds and hydrophobic contacts contributed to its strong binding affinity. Bacopasaponin D also exhibited a binding energy of -8.6 kcal/mol (Table 1), forming five hydrogen bonds with Asp32, Gly34, Asp228, Arg235 and Thr329 (Fig. 3e). Its extensive hydrophobic interactions with thirteen residues, including Ser35, Tyr71, and Ile226 (Table 2), anchored it firmly in the binding pocket, suggesting high inhibitory potential. Bacopaside 1 matched the binding energy of the other *Bacopa monnieri* compounds at -8.6 kcal/mol (Table 1), but formed the most extensive hydrogen bonding network among all compounds, with ten hydrogen bonds involving both catalytic aspartates (Asp32 and Asp228) and key residues like Gly34, Thr72, and Arg235 (Fig. 3f). While it had fewer hydrophobic interactions (six residues, Table 2), its robust hydrogen bonding network likely compensated for this, contributing to its high binding affinity. Furthermore, molecular docking performed using DockThor and CBDock indicated that these phytochemicals exhibit enhanced binding affinity for BACE1 compared to other phytochemicals (see Supplementary Table S1). The *Bacopa monnieri* phytochemicals demonstrated binding energies comparable to or better than the known BACE1 inhibitors. Notably, they formed more extensive hydrogen bonding networks, particularly Bacopaside 1, which could potentially make them promising candidates for developing novel BACE1 inhibitors. These findings suggest that phytochemicals from *Bacopa monnieri* could serve as valuable leads for Alzheimer’s disease treatment, offering a natural alternative to traditional inhibitors.

**Figure 3:**
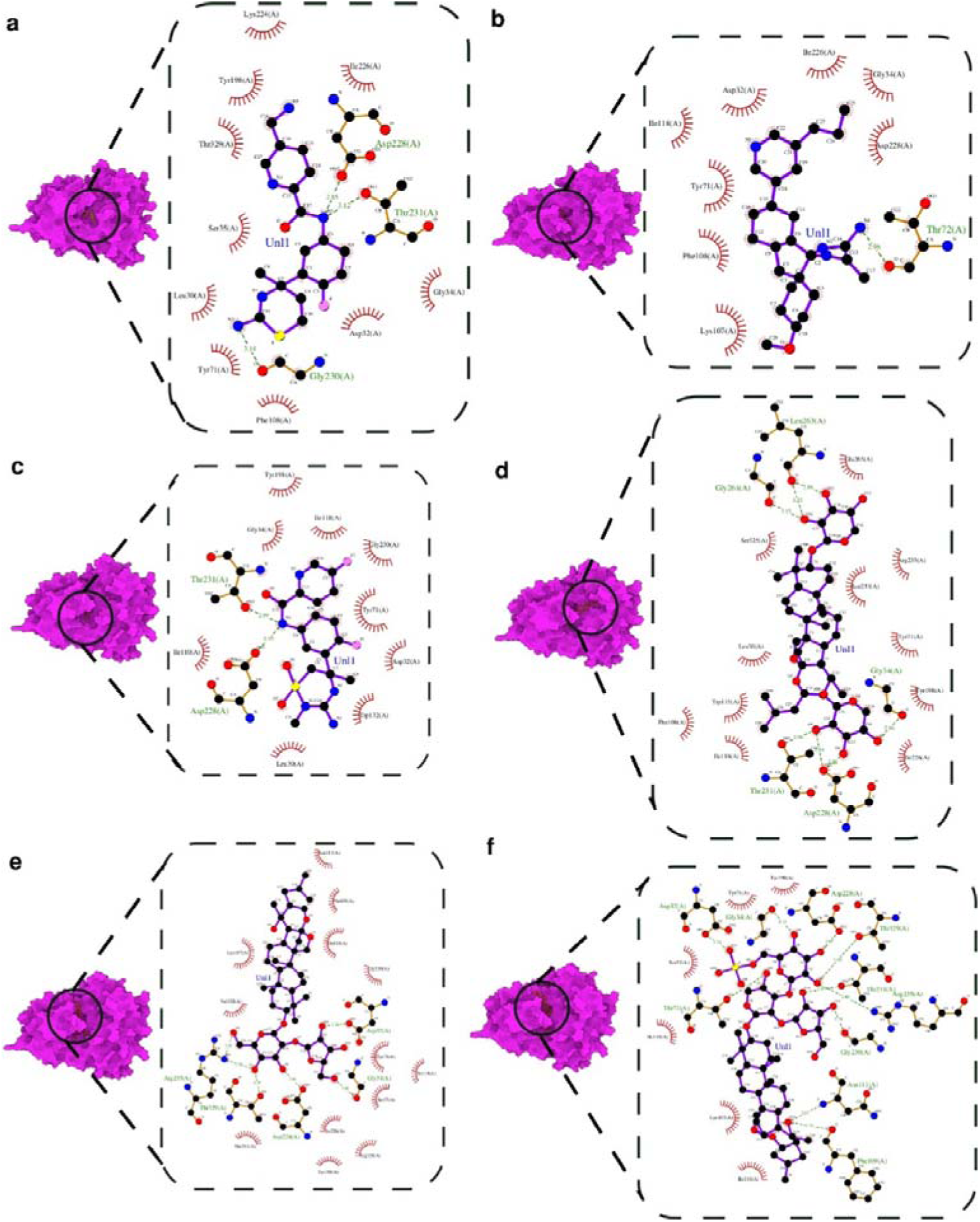
Molecular Docking interaction of BACE1 with (a) Atabecestat (b) Lanabecestat (c) Verubecestat (d) Bacopasaponin A (e) Bacopasaponin D (f) Bacopaside 1. BACE1 is shown in molecular surface view (magenta) and the interactions are shown in the zoomed in plots. Hydrogen bonding is shown in green colour and hydrophobic interaction are shown in red colour.

**Table 1:**
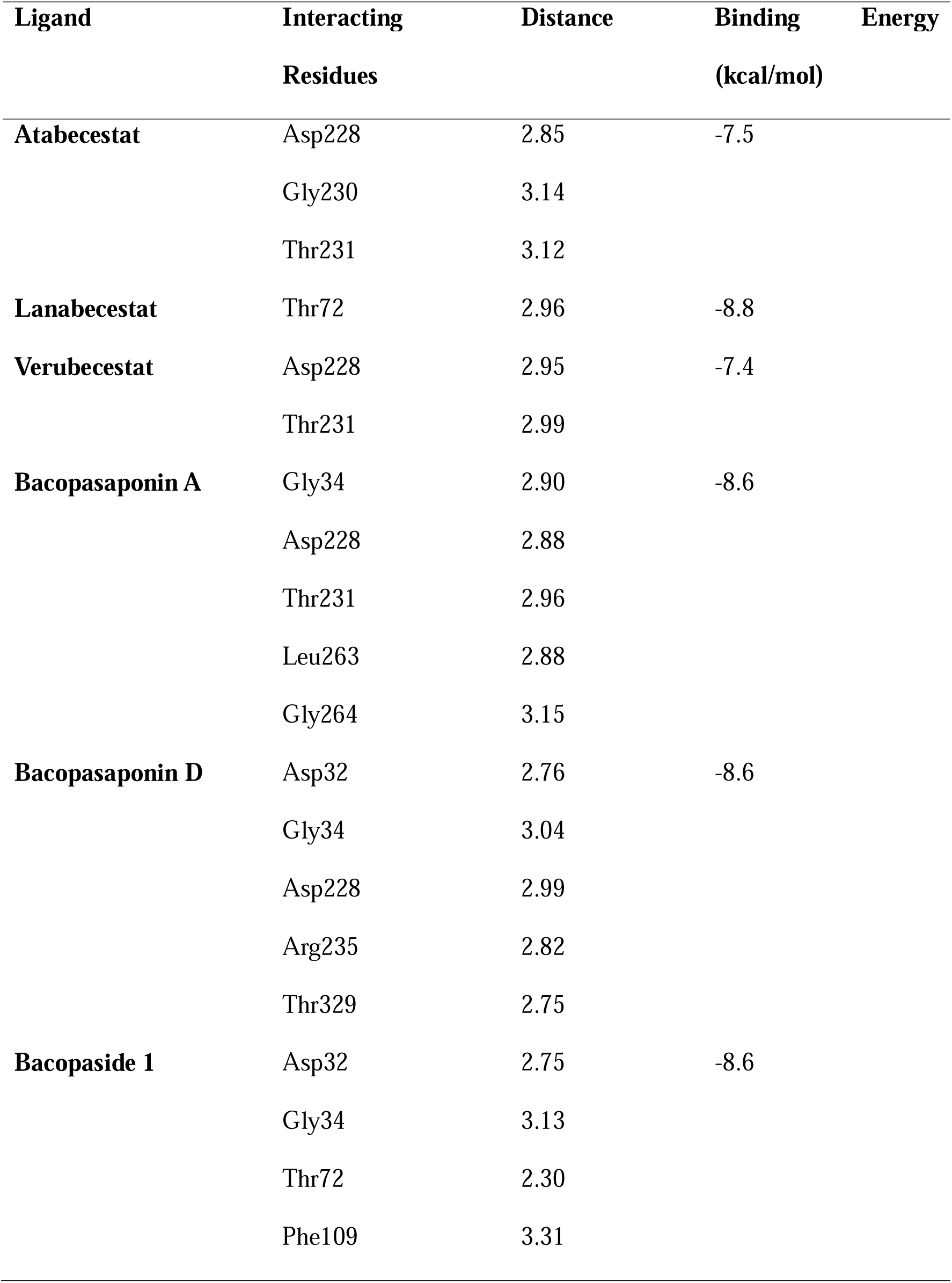
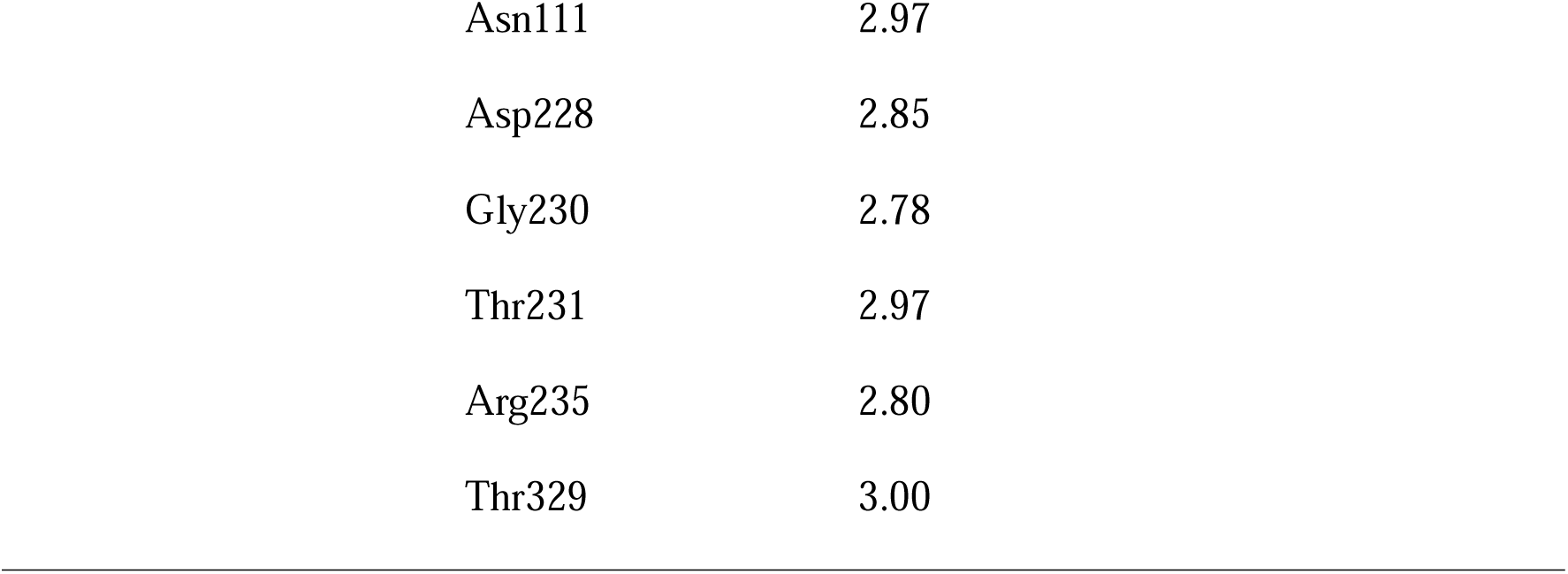
Molecular Docking details of residues of BACE1 interacting via Hydrogen Bonding.

**Table 2:** Molecular Docking details of residues of BACE1 interacting via Hydrogen Bonding with respective ligands.

| Ligand | Interacting Residue |
| --- | --- |
| Atabecestat | Leu30, Asp32, Gly34, Ser35, Tyr71, Phe108, Tyr198, Lys224, Ile226, Thr329 |
| Lanabecestat | Asp32, Gly34, Tyr71, Lys107, Phe108, Ile118, Ile226, Asp228, |
| Verubecestat | Leu30, Asp32, Gly34, Tyr71, Ile110, Ile118, Trp132, Tyr198, Gly230 |
| Bacopasaponin A | Leu30, Tyr71, Phe108, Trp115, Ile118, Tyr198, Ile226, Asn233, Arg235, Glu265, Ser325 |
| Bacopasaponin D | Ser35, Tyr71, Lys107, Phe109, Ile110, Ile118, Asn111, Arg128, Tyr198, Ile226, Gly230, Thr231, Val332 |
| Bacopaside 1 | Ser35, Tyr71, Lys107, Ile110, Ile118, Tyr198 |

**Table 3:**
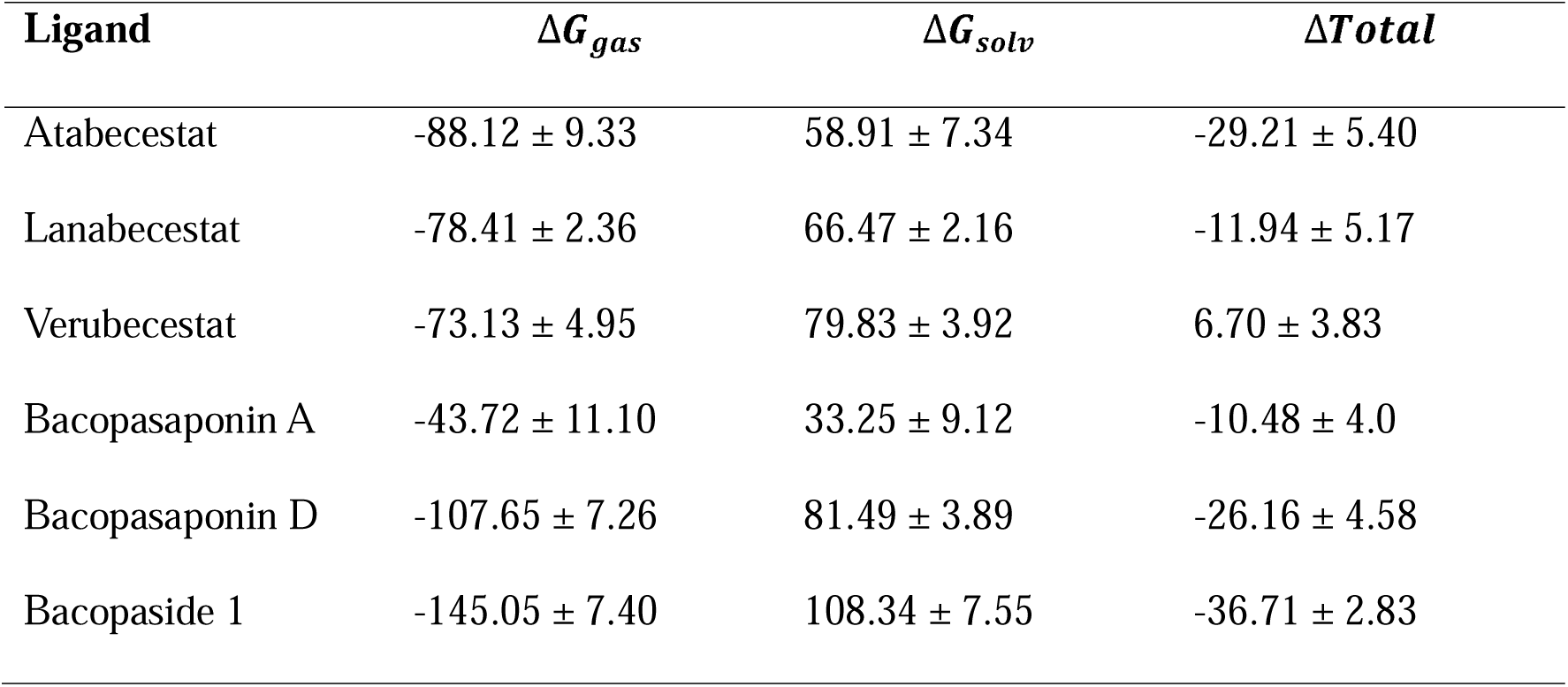
MMPBSA analysis of BACE1 in complex with different ligands.

### Molecular Dynamic Simulation

The BACE1 protein-ligand complexes were subjected to a 150 ns MD simulation. In order to account for potential sampling bias, we ran the MD Simulations three times. The details of the triplicate simulation are provided in the supplementary file (see Supplementary Fig. S2-S9).

### Root Mean Square Deviation (RMSD)

The Root Mean Square Deviation (RMSD) for BACE1 in complex with three known inhibitors and phytochemicals from *Bacopa monnieri* was analysed over a 150 ns molecular dynamics (MD) simulation to assess stability. RMSD values for all complexes were generally stable, ranging between 0.1 and 0.3 nm, with an initial adjustment phase observed during the first 20 ns. After equilibration, the RMSD profiles stabilized, indicating that the complexes reached a steady conformation. Among the inhibitors, Verubecestat exhibited the highest RMSD values, slightly exceeding 0.3 nm, suggesting greater conformational flexibility (Fig. 4a, blue). Lanabecestat (Fig. 4a, green) and Atabecestat (Fig. 4a, red) displayed more conservative profiles, maintaining RMSD values within 0.2 to 0.3 nm, indicating similar stabilization of BACE1. The *Bacopa monnieri* phytochemicals demonstrated promising stability. Bacopaside I closely matched Lanabecestat, with RMSD values between 0.2 and 0.25 nm (Fig. 4a, maroon), suggesting it induces similar stability in BACE1. Bacopasaponin A (Fig. 4a, magenta) and Bacopasaponin D (Fig. 4a, cyan) also showed stable profiles, with RMSD values ranging from 0.2 to 0.3 nm. Bacopasaponin D exhibited slightly higher flexibility, akin to Verubecestat, potentially indicating dynamic binding behaviour. Overall, the phytochemicals, particularly Bacopaside I, exhibited RMSD profiles comparable to the known inhibitors, suggesting they stabilize BACE1 similarly without introducing excessive conformational strain, indicating potential for effective inhibition.

**Figure 4:**
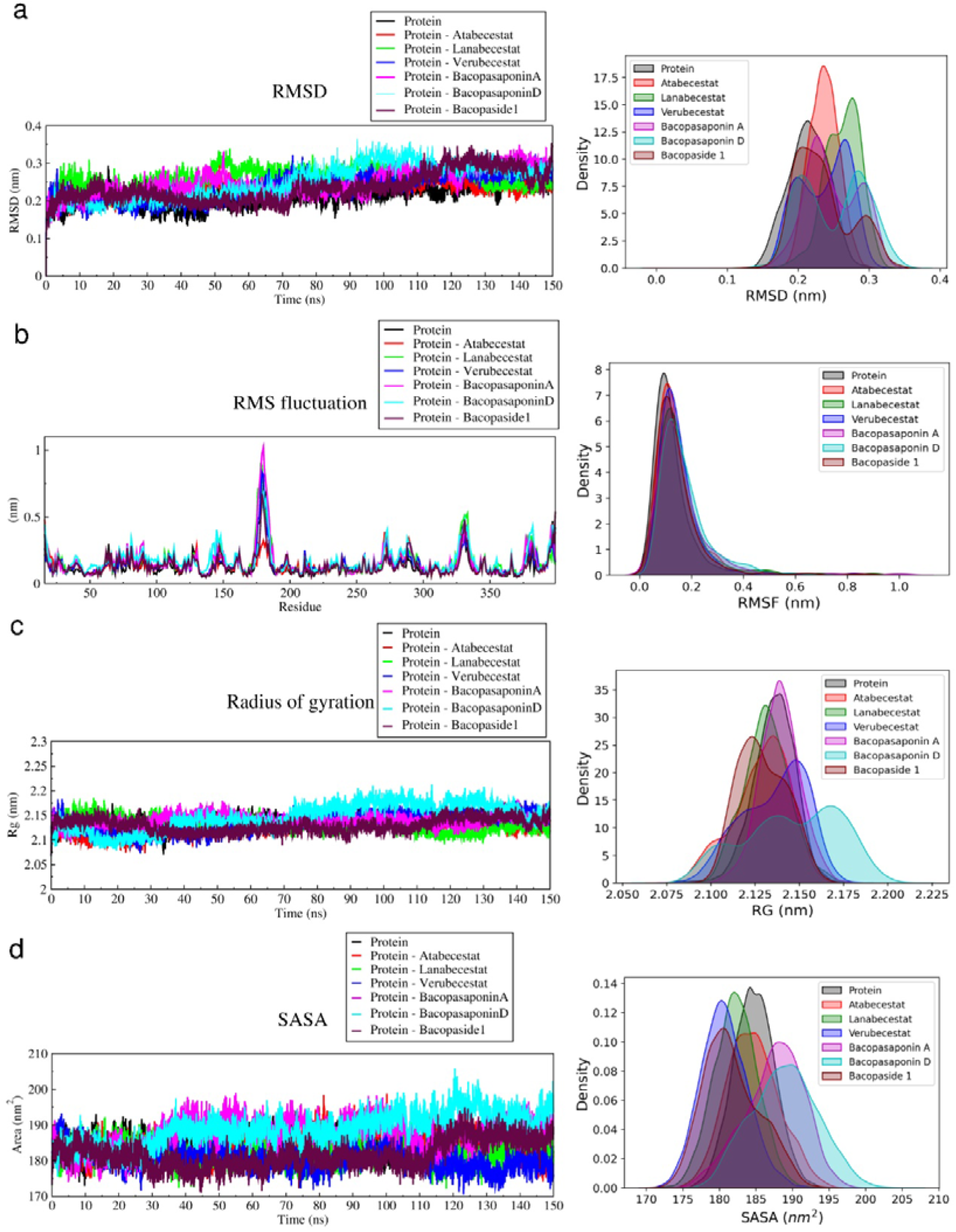
150 ns Molecular Dynamic Simulation analysis of BACE1 showing (a) RMSD (b) RMSF (c) Radius of Gyration and (d) SASA analysis in apo form (black) and in association with Atabecestat (red), Lanabecestat (green), Verubecestat (blue), Bacopasaponin A (magenta), Bacopasaponin D (cyan) and Bacopaside 1 (maroon). Each analysis has their corresponding probability distribution curve with the same colour code.

### Root Mean Square Fluctuation (RMSF)

The Root Mean Square Fluctuation (RMSF) analysis was performed to assess the flexibility of specific BACE1 residues bound to different ligands, providing insights into their binding dynamics and stability. Overall, BACE1 exhibited low RMSF values, indicating structural stability across ligand-bound states. Key active site residues, Asp32 and Asp228, remained highly stable (RMSF < 0.2 nm) in all cases, confirming that the ligands effectively stabilize the catalytic core, preserving BACE1’s enzymatic function. The flap region, crucial for substrate binding, displayed moderate flexibility (RMSF ∼0.2-0.3 nm) with ligand-specific variations. Bacopasaponin A induced the highest flap flexibility (RMSF ∼0.4 nm), suggesting a unique inhibitory mechanism through flap modulation (Fig. 4b, magenta). Atabecestat (Fig. 4b, red) and Verubecestat (Fig. 4b, blue) showed moderate effects, while Lanabecestat (Fig. 4b, green), Bacopaside I (Fig. 4b, maroon), and Bacopasaponin D (Fig. 4b, cyan) resulted in lower flap movement, indicating tighter binding. The N-terminal 10S loop exhibited high flexibility (RMSF > 0.3 nm) in all ligand-bound states, largely independent of ligand binding, potentially playing a role in substrate recognition or product release. Bacopasaponin D and Lanabecestat induced slightly higher fluctuations in this region. Similarly, the 113S loop displayed moderate flexibility (RMSF ∼0.2-0.3 nm) with ligand-specific effects. Bacopasaponin A and Bacopaside I caused higher flexibility (∼0.4 nm), which might impact substrate access to the active site, while the known inhibitors (Atabecestat, Lanabecestat, Verubecestat) showed moderate effects. A hyper-flexible region near Insert A (residues 175-185) exhibited the highest flexibility among all regions (RMSF > 0.8 nm), with notable ligand-dependent variations. Bacopasaponin A induced the most significant fluctuations (RMSF > 1.0 nm), indicating a unique interaction mode that might alter BACE1’s overall conformation. Lanabecestat and Verubecestat had moderate effects (∼0.8 nm), while Atabecestat, Bacopasaponin D, and Bacopaside I induced lower fluctuations. Insert D (residues 270-274), a short loop region, showed moderate flexibility (∼0.2-0.3 nm) with distinct ligand effects. Bacopasaponin D and Bacopasaponin A induced the highest flexibility (∼0.4 nm), suggesting potential interactions that may affect BACE1’s C-terminal domain. Atabecestat and Verubecestat caused moderate effects, while Lanabecestat and Bacopaside I led to minimal movement, possibly stabilizing this region of the protein. Finally, Insert F (residues 311-317) exhibited similar flexibility to Insert D, with RMSF values ranging from 0.05 to 0.1 nm across ligands. Bacopasaponin D caused the most significant fluctuations (∼0.1 nm), indicating a unique interaction mode that may influence the conformation of BACE1’s C-terminal domain. Lanabecestat, Verubecestat, and Bacopasaponin A showed moderate effects (∼0.15 nm), while Atabecestat and Bacopaside I induced lower fluctuations (∼0.1 nm). The RMSF analysis indicates that *Bacopa monnieri* phytochemicals, especially Bacopaside I, show stability and flexibility profiles comparable to known inhibitors, particularly Atabecestat and Lanabecestat. The slightly higher flexibility observed with Bacopasaponin A and Bacopasaponin D may reflect a more adaptable binding mode, which could be advantageous for targeting different conformations of BACE1 but might also introduce some instability. Bacopaside I’s RMSF profile aligns closely with those of the synthetic inhibitors, suggesting it may stabilize key protein regions while potentially avoiding the issues that led to the clinical failure of Lanabecestat and Atabecestat.

### Radius of Gyration (Rg)

The Radius of Gyration (Rg) measures the compactness of a protein, providing insights into its stability and folding during a molecular dynamics (MD) simulation. The unbound BACE1 protein (Fig. 4c, black) fluctuates around 2.1 nm, indicating a stable, compact structure. This serves as a baseline for evaluating ligand effects. Atabecestat (Fig. 4c, red) and Lanabecestat (Fig. 4c, green) closely mirror the unbound state, with Rg values around 2.1 nm, indicating that these ligands stabilize the protein without significant structural changes. Verubecestat (Fig. 4c, blue) shows slightly higher fluctuations, occasionally reaching 2.15 nm, suggesting minor conformational changes that slightly reduce compactness but still maintain overall stability. Bacopasaponin A (Fig. 4c, magenta) and Bacopasaponin D (Fig. 4c, cyan) also show similar Rg profiles to the known inhibitors. However, Bacopasaponin D exhibits a slight increase in Rg after 70 ns, reaching up to 2.25 nm, indicating potential expansion and a more flexible interaction. Bacopaside I (Fig. 4c, maroon) maintains a stable Rg around 2.1 nm, suggesting that it stabilizes the protein similarly to Atabecestat and Lanabecestat. Overall, Rg values for BACE1-ligand complexes range from 2.05 to 2.25 nm. The slight increases for Verubecestat and Bacopasaponin D suggest some flexibility in binding, while Atabecestat, Lanabecestat, and Bacopaside I stabilize the protein in a compact conformation. This stability is likely beneficial for inhibiting BACE1 activity, as it may prevent excessive conformational changes that could affect the enzyme’s function. Bacopaside I’s stability profile makes it a promising candidate for further study.

### Solvent Accessible Surface Area (SASA)

The Solvent Accessible Surface Area (SASA) simulation over 150 nanoseconds provides key insights into the dynamic interactions between BACE1 and its ligands, including both known inhibitors and phytochemicals. The SASA plot reveals time-dependent fluctuations for all compounds, highlighting dynamic protein-ligand interactions. Notably, the phytochemicals display SASA profiles similar to the clinically-tested inhibitors, suggesting that they may bind to BACE1 in a comparable manner, potentially offering similar inhibitory effects. Among the phytochemicals, Bacopasaponin D (Fig. 4d, cyan) exhibits the highest average SASA values (around 190-200 nm²) and the most significant fluctuations, paralleling Atabecestat’s behaviour. These elevated values suggest extensive interactions with BACE1’s surface, possibly due to Bacopasaponin D’s structural complexity. This dynamic interaction may contribute to a stable binding mode, enhancing its potential as an effective inhibitor. Bacopasaponin A (Fig. 4d, magenta) and Bacopaside I (Fig. 4d, maroon) display SASA profiles similar to Verubecestat (Fig. 4d, blue) and Lanabecestat (Fig. 4d, green) (approximately 180-190 nm²), indicating that these phytochemicals may occupy a similar volume within the BACE1 binding site. Their consistent SASA values throughout the simulation suggest stable interactions with BACE1, a trait often linked with effective inhibitors. The periodic fluctuations in SASA values for all compounds, including both inhibitors and phytochemicals, reflect conformational changes in the protein-ligand complexes. This suggests that the phytochemicals induce structural adaptations in BACE1 akin to those triggered by the known inhibitors, contributing to their inhibitory potential. Bacopasaponin D and Atabecestat show synchronized increases in SASA values and fluctuations after 90 ns, indicating that both compounds may induce large-scale conformational changes in BACE1 over time. This sustained effect could signify prolonged inhibitory action, an advantageous feature for therapeutic agents. Bacopasaponin A and Bacopaside I maintain stable SASA ranges, mirroring Verubecestat and Lanabecestat’s behaviour, indicating robust interactions with BACE1. Slightly lower SASA values for Bacopaside I suggest a more compact binding mode, possibly leading to higher binding affinity. While synthetic inhibitors have faced challenges in clinical trials, the similar SASA profiles of Bacopa monnieri phytochemicals suggest that these natural compounds may offer improved safety and efficacy, addressing limitations faced by synthetic drugs.

### Molecular Mechanics Poisson-Boltzmann Surface Area (MMPBSA)

The MMPBSA (Molecular Mechanics Poisson-Boltzmann Surface Area) analysis reveals binding free energies (ΔG) of various ligands interacting with BACE1, offering key insights into their binding affinities^31^. Lower (more negative) ΔG values represent stronger binding. Among the clinically tested inhibitors, Atabecestat has the most favourable ΔG (-29.21 kcal/mol) (Fig. 5), while Lanabecestat (Fig. 5) shows a moderately favourable binding energy (-11.94 kcal/mol). Interestingly, Verubecestat has a positive ΔG value (+6.7 kcal/mol) (Fig. 5), indicating unfavourable binding under these conditions. The *Bacopa monnieri* phytochemicals demonstrate highly favourable binding energies, surpassing or matching those of the clinically tested inhibitors. Bacopaside 1 shows the most negative ΔG (-36.71 kcal/mol) (Fig. 5), suggesting the strongest binding affinity among all tested compounds. Bacopasaponin D also exhibits a strong ΔG (-26.16 kcal/mol) (Fig. 5), comparable to Atabecestat. Bacopasaponin A, with a ΔG of -10.48 kcal/mol (Fig. 5), shows a binding affinity similar to Lanabecestat. These findings suggest that *Bacopa monnieri* phytochemicals, especially Bacopaside 1 and Bacopasaponin D, may overcome some limitations faced by synthetic inhibitors in clinical trials. The strong binding affinities of these phytochemicals indicate that they could form stable interactions with BACE1, potentially leading to effective enzyme inhibition. The variability in ΔG values for clinically tested inhibitors, ranging from highly favourable for Atabecestat to unfavourable for Verubecestat, underscores the complexity of translating binding affinity to clinical efficacy. Notably, Bacopaside 1’s exceptionally low ΔG suggests it forms multiple strong interactions within the BACE1 binding site. The phytochemicals, particularly Bacopaside 1 and Bacopasaponin D, show small error bars, indicating consistent binding modes across the simulation. Overall, the MMPBSA results align with SASA findings, further supporting the potential of *Bacopa monnieri* phytochemicals as BACE1 inhibitors, with Bacopaside 1 being particularly promising for future drug development.

**Figure 5:**
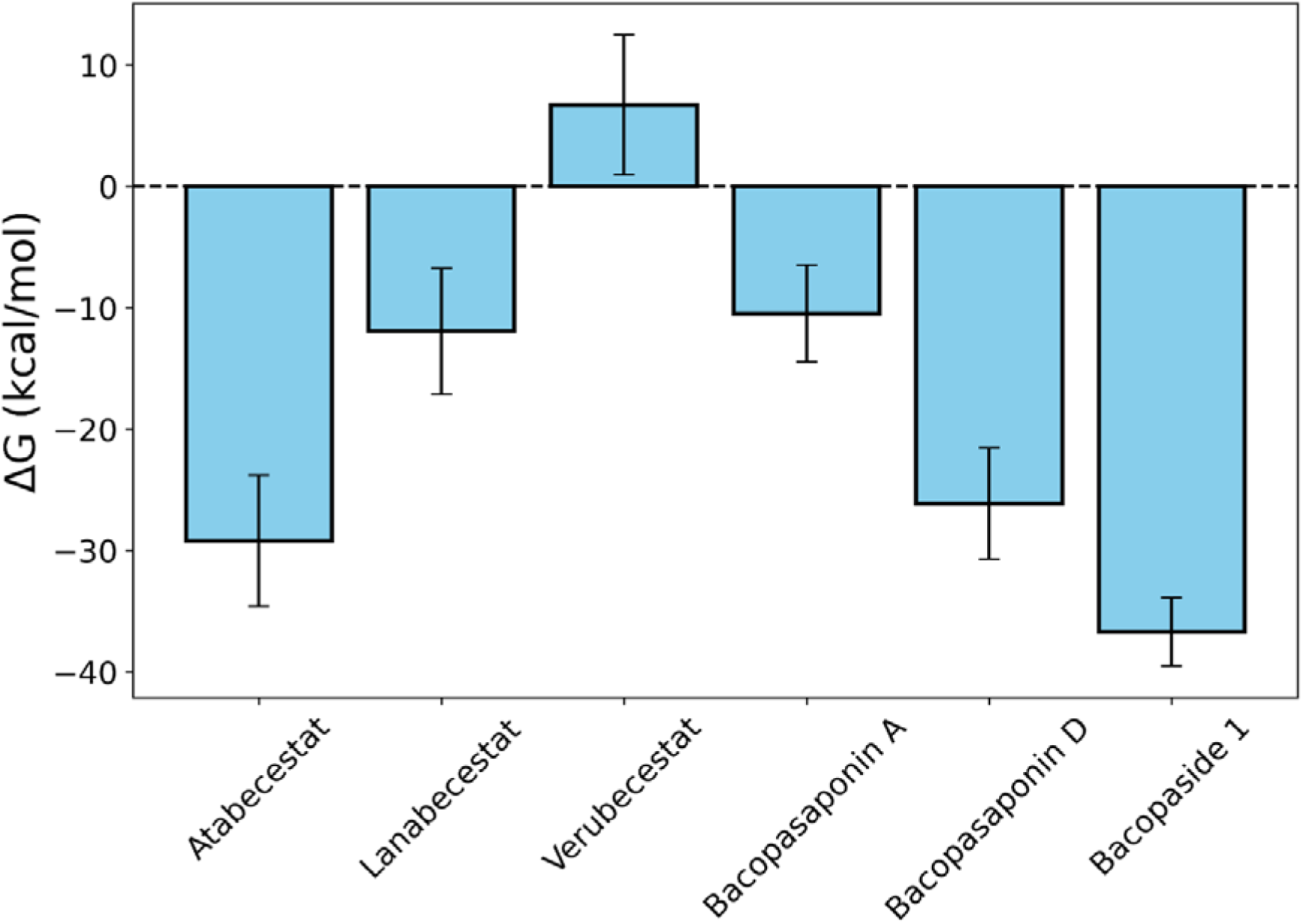
MM-PBSA analysis of BACE1 with respective ligands. The error bar denotes the standard deviation associated with each ligand’s energy

### Dynamic Cross Correlation Matrix (DCCM) Analysis

The Dynamic Cross-Correlation Matrix (DCCM) analysis provides crucial insights into the correlated motions of residues within the BACE1 active site when bound to different ligands. This comprehensive examination reveals intriguing patterns of residue interactions that may explain their potential effectiveness as BACE1 inhibitors. The DCCM heatmaps showcase a range of correlations from strong positive (red, +1) to strong negative (blue, -1) between residues 225-235 and 25-35, which encompass the critical catalytic dyad (residues 32 and 228) of BACE1. This region is pivotal for the enzyme’s function, and alterations in its dynamic behaviour can significantly impact BACE1’s catalytic activity. Atabecestat (Fig. 6a) exhibits a moderate level of anticorrelation across the matrix, particularly between residues 27-31 and 229-233. This suggests a coordinated movement that may stabilize the binding pocket, potentially explaining its effectiveness in vitro. Lanabecestat (Fig. 6b) shows a similar pattern to Atabecestat but with slightly weaker anticorrelations. The consistency in pattern between these two compounds might indicate a common mechanism of action. Verubecestat (Fig. 6c) presents a notably different profile, with weaker anticorrelations and some positive correlations, especially around residues 32-34. This distinct pattern could be linked to its different clinical outcomes compared to Atabecestat and Lanabecestat. These mixed correlations suggest that known inhibitors may only partially disrupt the catalytic dyad formed by Asp32 and Asp228, allowing some degree of coordinated motion between these residues. Intriguingly, the *Bacopa monnieri* phytochemicals demonstrate DCCM profiles that not only mimic but in some aspects surpass those of the clinically tested inhibitors. Bacopasaponin A (Fig. 6d) shows a striking similarity to Atabecestat’s profile, with strong anticorrelations between residues 27-31 and 229-233. This remarkable resemblance suggests that Bacopasaponin A might induce similar conformational changes in BACE1, potentially leading to comparable inhibitory effects. Bacopasaponin D (Fig. 6e) exhibits the strongest and most extensive anticorrelations among all compounds. The intense blue regions, particularly between residues 26-31 and 230-234, indicate highly coordinated opposite motions. This strong anticorrelation near the catalytic dyad (32 and 228) suggests that Bacopasaponin D might induce substantial conformational changes, potentially leading to more effective enzyme inhibition. Bacopaside 1 (Fig. 6f) presents a unique profile with moderate anticorrelations spread more evenly across the matrix. Notably, it shows distinct patterns around residues 32 and 228, suggesting a specific interaction with the catalytic dyad that differs from the other compounds.

**Figure 6:**
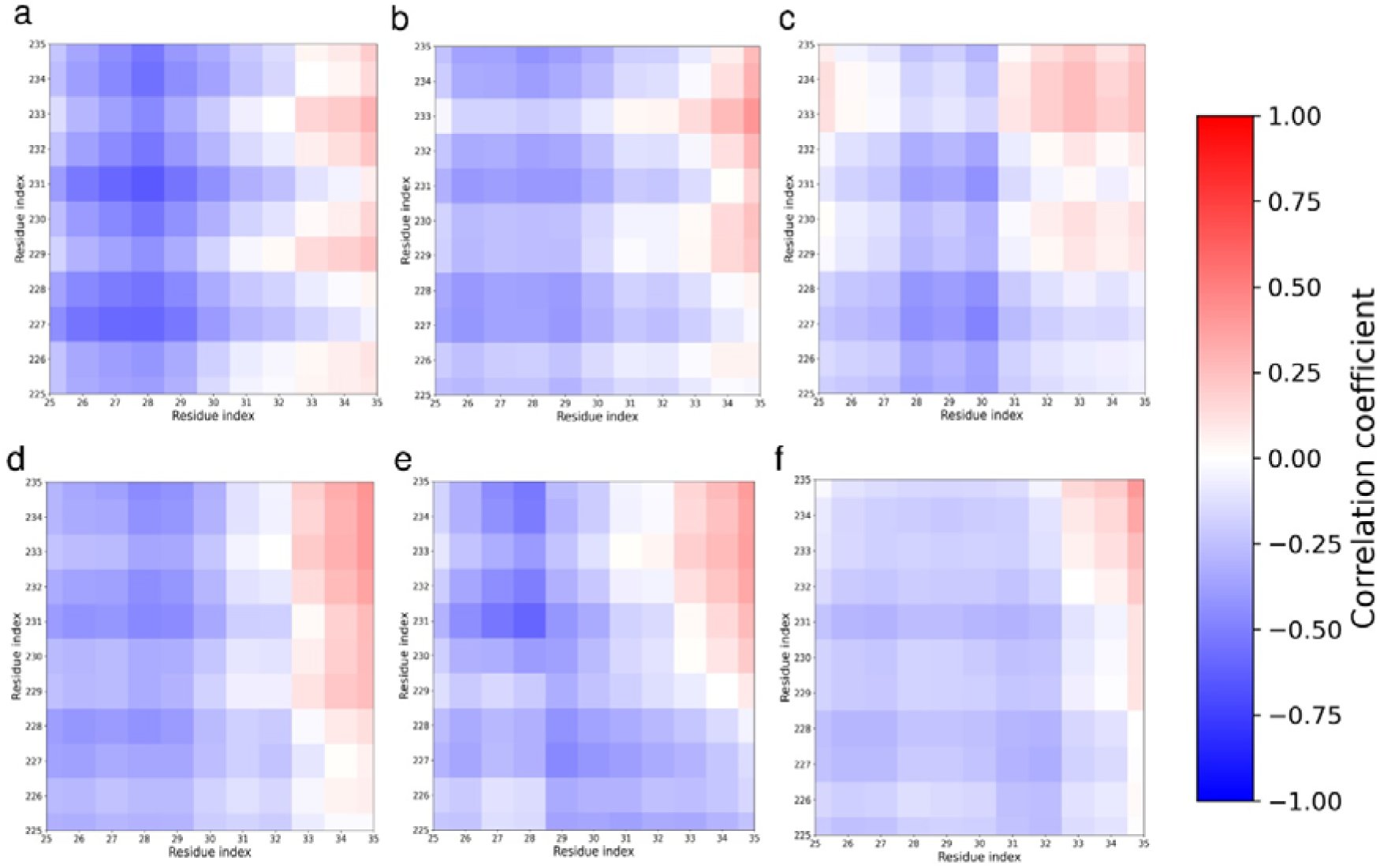
Dynamic Cross Correlation of BACE1 active site 1 (residue 25-35, including Asp32) and active site 2 (residues 225-235, including Asp228) with (a) Atabecestat (b) Lanabecestat (c) Verubecestat (d) Bacopasaponin A (e) Bacopasaponin D (f) Bacopaside 1

Moreover, particularly in their interactions with key regions like the 10s loop (residues 5-16) (Fig. 7) and the 113s loop (residues 105-117) (Fig. S11), known inhibitors show varied responses to the 10s loop. For active site 1 (Asp32), these inhibitors display mixed or weak positive correlations with the upper portion of the 10s loop (residues 9-16) (Fig. 7a-c), indicating partial flexibility in this region. In contrast, their interactions with active site 2 (Asp228) are characterized by strong anti-correlations in the upper 10s loop (Fig. 8a-c), suggesting a decoupling effect that may hinder dynamic coordination between this loop and the catalytic dyad. The inhibitors also induce complex correlation patterns with the 113s loop, which is essential for substrate binding and catalysis. Atabecestat exhibits mixed correlations, with negative correlations in the lower 113s loop (residues 105-110) and positive correlations with residues 28-32 in the upper loop (Fig. S11a). Lanabecestat (Fig. S11b) and Verubecestat (Fig. S11c) follow similar trends but with stronger negative correlations across most of the 113s loop, pointing to selective stabilization of specific regions of the loop, which may contribute to a partially rigid conformation, limiting substrate access. In contrast, the phytochemicals exhibit stronger and more consistent positive correlations between the 10s loop and active site 1 (Fig. 7d-f), particularly Bacopaside I, which shows uniform positive correlations with weak negative correlation in upper and lower residues. This suggests enhanced coordination and flexibility between the 10s loop and the catalytic residue Asp32, potentially improving substrate adaptability.

**Figure 7:**
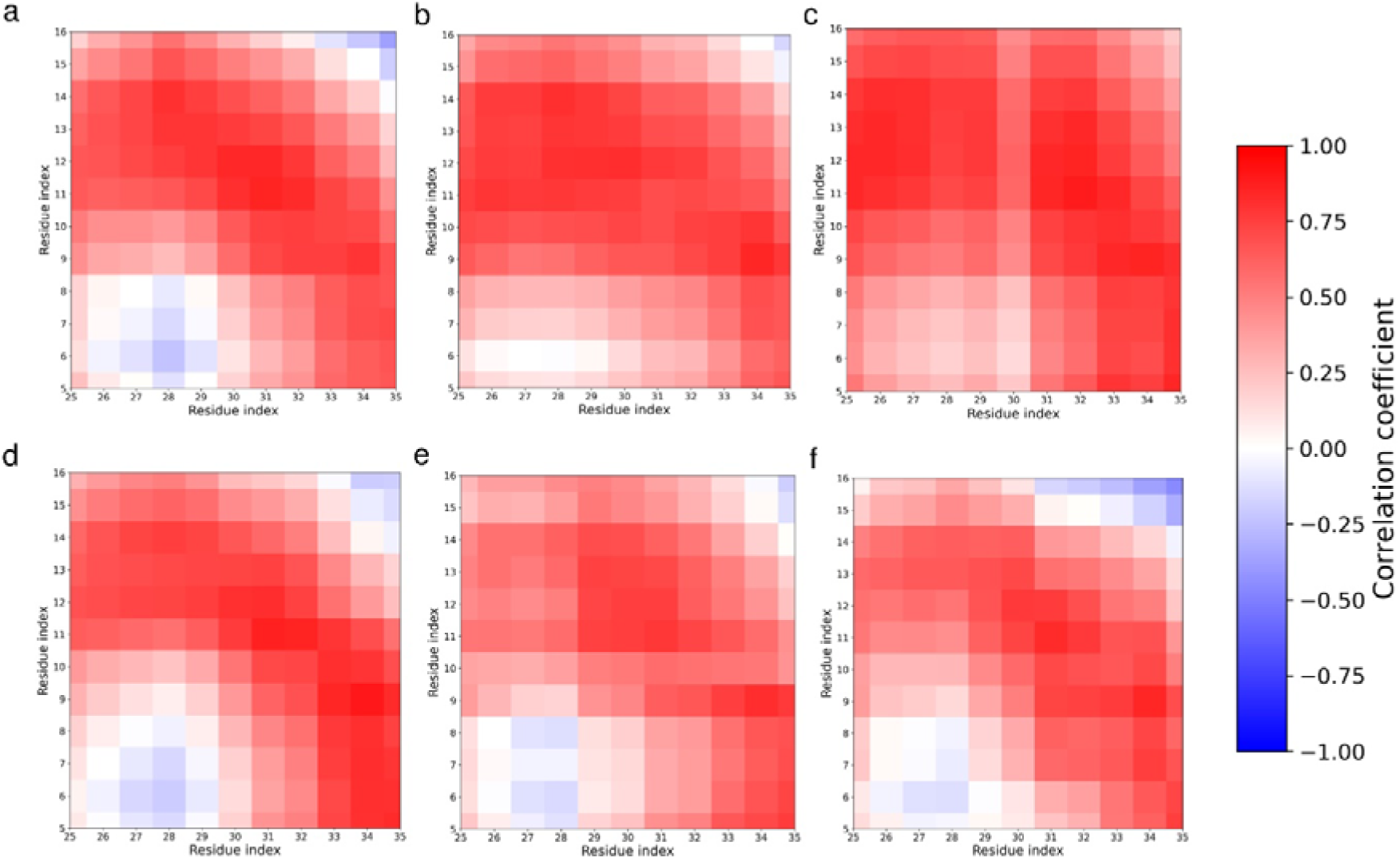
Dynamic Cross Correlation of BACE1 active site 1 (residue 25-35, including Asp32) and 10S loop region (residues 5-16) with (a) Atabecestat (b) Lanabecestat (c) Verubecestat (d) Bacopasaponin A (e) Bacopasaponin D (f) Bacopaside 1

**Figure 8:**
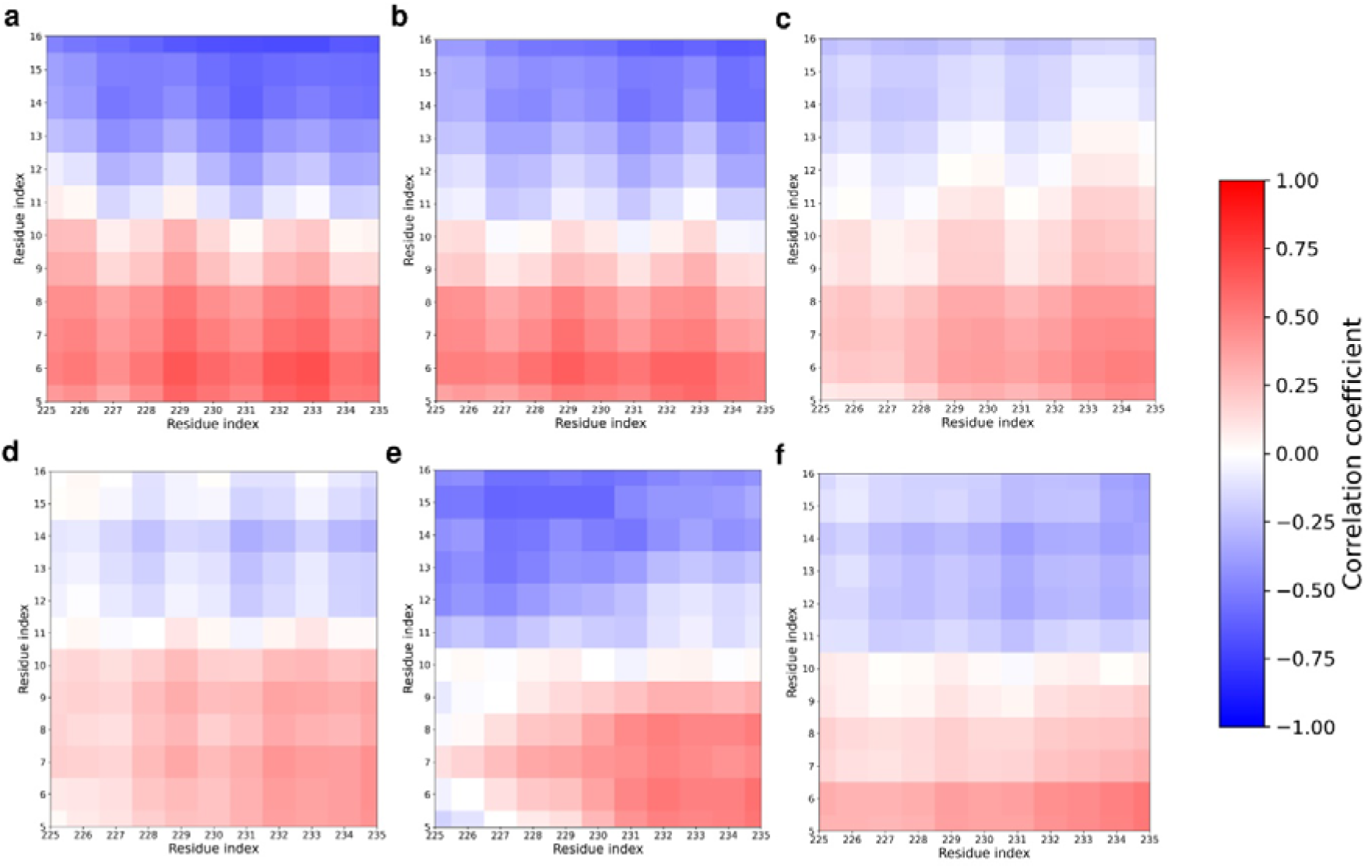
Dynamic Cross Correlation of BACE1 active site 2 (residue 225-235, including Asp32) and 10S loop region (residues 5-16) with (a) Atabecestat (b) Lanabecestat (c) Verubecestat (d) Bacopasaponin A (e) Bacopasaponin D (f) Bacopaside 1

Moreover, phytochemicals induce broader anti-correlations between the 10s loop and active site 2 (Fig. 8d-f), indicating more extensive disruption of enzyme dynamics. The 113s loop responses are also strikingly different. Bacopaside, I induce strong and uniform negative correlations across the 113s loop (Fig. S11f), suggesting that the phytochemicals promote a more dynamic and flexible 113s loop conformation, potentially disrupting its ability to properly align the substrate for catalysis. These dynamic modulations may allow phytochemicals to achieve a more nuanced inhibitory mechanism, enhancing flexibility and disrupting enzyme function more effectively than the rigid active site blockade imposed by synthetic inhibitors. This differential regulation of BACE1 dynamics highlights the potential of *Bacopa monnieri* phytochemicals as superior BACE1 inhibitors, offering more refined control of enzyme conformation and improved therapeutic prospects. The DCCM analysis suggests that phytochemicals from *Bacopa monnieri*, particularly Bacopaside 1, induce more pronounced and disruptive effects on BACE1’s catalytic dyad and active site dynamics than known inhibitors. These findings offer valuable insights into alternative mechanisms of BACE1 inhibition and underscore the potential of natural compounds as effective therapeutic agents for neurodegenerative diseases.

### Deformity Analysis

The deformity analysis of BACE1 residues upon ligand binding offers valuable insights into the structural dynamics of enzyme-inhibitor complexes, shedding light on their mechanisms of action and potential efficacy as inhibitors. Atabecestat induces significant deformations across the BACE1 structure (Fig. 9a), especially around residues 70-80 and 250-300, likely corresponding to flexible regions important for enzyme function. The active site residues show moderate deformation, indicating a direct interaction that contributes to its inhibitory effect. Lanabecestat exhibits a similar deformation pattern but with lower magnitudes (Fig. 9b), which may explain its different clinical profile, suggesting a less disruptive binding mode. Verubecestat presents a unique deformation profile (Fig. 9c), with pronounced changes around residues 70-80 and 170-180, likely affecting regions critical for substrate recognition or catalysis. The *Bacopa monnieri* phytochemicals exhibit notably different deformation profiles. Bacopasaponin A induces minimal deformation across the BACE1 structure (Fig. 9d), suggesting it stabilizes the enzyme in an unfavourable conformation for catalysis without causing significant structural changes. Bacopasaponin D displays moderate deformations around specific regions (residues 150-180) (Fig. 9e), indicating a more selective interaction with key functional areas, potentially offering a targeted inhibitory mechanism. Bacopaside 1 induces the least deformation (Fig. 9f), suggesting it stabilizes BACE1 in an inactive conformation, offering a more consistent inhibitory mechanism. The stabilization of the active site region (residues 32 and 228) by the phytochemicals likely prevents conformational changes necessary for BACE1’s catalytic activity, a more effective strategy than that of the clinical compounds, which show higher deformation in these areas. Bacopasaponin D’s selective deformation may indicate specific, strong interactions that contribute to higher specificity and fewer off-target effects. The varied deformation profiles among the phytochemicals suggest different mechanisms for inhibiting BACE1, which could be beneficial in addressing diverse patient responses or overcoming resistance. These findings highlight the potential of these phytochemicals as promising candidates for BACE1 inhibition and drug development.

**Figure 9:**
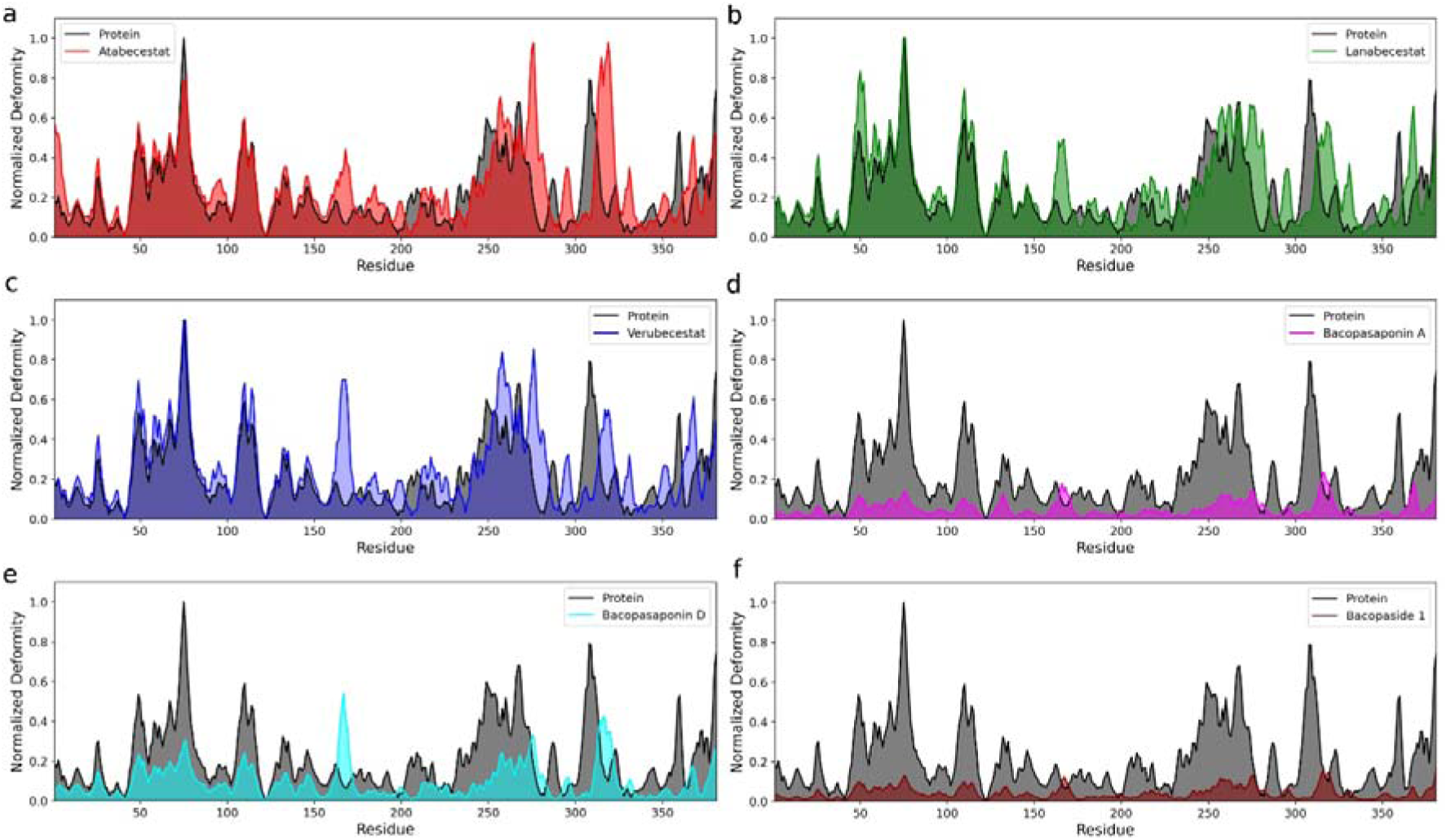
Deformity analysis of BACE1 with (a) Atabecestat (b) Lanabecestat (c) Verubecestat (d) Bacopasaponin A (e) Bacopasaponin D and (f) Bacopaside 1. The black curve is the deformity of the apo form of the protein.

### Principal Component Analysis

The Principal Component Analysis (PCA) of BACE1 complexed with various ligands provides valuable insights into the dynamic behaviour of these complexes, highlighting the effects of each ligand on protein flexibility and stability. The PCA plots, which show the first three principal components (PC1, PC2, and PC3) against residue positions, reveal distinct patterns of protein dynamics induced by different ligands. For Atabecestat, PC1 shows a decreasing trend across residues (Fig. 10a, black), indicating reduced global motion and stabilization of BACE1, especially in the C-terminal region. The fluctuations in PC2 (Fig. 10a, red) and PC3 (Fig. 10a, blue) are moderate with notable peaks around residues 100-150 and 250-300, suggesting localized flexibility within flexible loops or secondary structures. This implies that Atabecestat stabilizes BACE1 while allowing necessary localized movements, which could facilitate enzyme activity. Lanabecestat displays a more uniform trend in PC1 (Fig. 10b, black), with less pronounced fluctuations in the protein’s initial regions, suggesting a more consistent global stabilization. However, PC2 (Fig. 10b, red) and PC3 (Fig. 10b, blue) show significant fluctuations, particularly towards the C-terminal, indicating that Lanabecestat provides strong global stabilization while allowing functional flexibility in specific regions. Verubecestat’s PCA results show a moderate decline in PC1 fluctuations (Fig. 10c, black), suggesting balanced stabilization. The slightly more pronounced fluctuations in PC2 (Fig. 10c, red) and PC3 (Fig. 10c, blue) compared to Atabecestat indicate that Verubecestat achieves both global and localized stabilization, supporting necessary conformational changes for enzyme function. Bacopasaponin A exhibits a trend similar to known inhibitors, with a gradual decrease in PC1 fluctuations (Fig. 10d, black), indicating effective global stabilization. The moderate fluctuations in PC2 (Fig. 10d, red) and PC3 (Fig. 10d, blue), with peaks similar to those of Lanabecestat and Verubecestat, suggest that Bacopasaponin A stabilizes BACE1 while maintaining necessary localized flexibility. Bacopasaponin D shows a stabilization pattern in PC1 (Fig. 10e, black) comparable to other phytochemicals and inhibitors. Its PC2 (Fig. 10e, red) and PC3 (Fig. 10e, blue) fluctuations, consistent with Lanabecestat and Verubecestat, indicate effective modulation of protein dynamics, particularly in the C-terminal region, similar to established inhibitors. Bacopaside I present a stabilization pattern in PC1 (Fig. 10f, black) like Bacopasaponin A and D, with pronounced localized flexibility in PC2 (Fig. 10f, red) and PC3 (Fig. 10f, blue), similar to known inhibitors. This suggests Bacopaside I, like other phytochemicals, maintains functional dynamics while providing stabilization. Overall, the PCA results demonstrate that Bacopasaponin A, Bacopasaponin D, and Bacopaside I exhibit stabilization patterns comparable to or potentially better than the known BACE1 inhibitors Atabecestat, Lanabecestat, and Verubecestat. The phytochemicals’ ability to stabilize BACE1 globally while preserving necessary localized motions suggests they could be effective inhibitors, possibly enhancing binding affinity and inhibitory potency.

**Figure 10:**
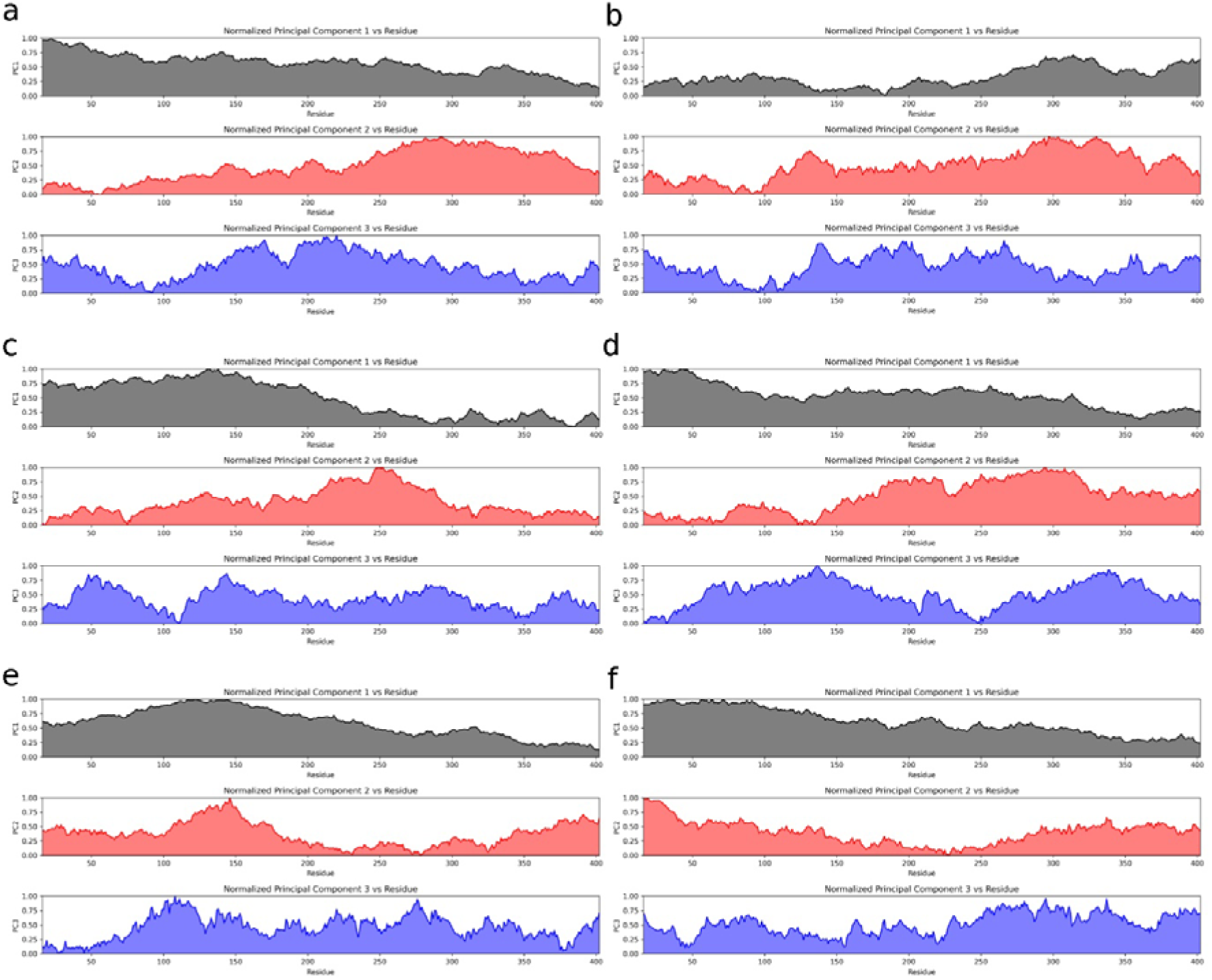
Principal Component Analysis (PCA) of BACE1 representing three distinct PCs: PC1 (black), PC2 (red), and PC3 (red) with (a) Atabecestat (b) Lanabecestat (c) Verubecestat (d) Bacopasaponin A (e) Bacopasaponin D and (f) Bacopaside 1 Moreover, the free energy landscape in the PCA space of BACE1 in complex with synthetic inhibitors and *Bacopa monnieri* phytochemicals revealed distinct free energy landscapes, providing crucial insights into their binding mechanisms and efficacy (Fig. S20). Synthetic inhibitors exhibited varied energy profiles. Atabecestat (Fig. S20a) and Verubecestat (Fig. S20c) showed broad distributions with multiple minima, indicating diverse but potentially unstable BACE1 conformations. In contrast, Lanabecestat (Fig. S20b) induced a well-defined, narrow energy basin, suggesting a highly stable, specific binding mode. Remarkably, *Bacopa monnieri* phytochemicals demonstrated more favourable energy landscapes. Bacopasaponin A (Fig. S20d) and Bacopasaponin D (Fig. S20e) exhibited confined yet diverse low-energy states, indicating a balance between flexibility and stability. Notably, Bacopaside 1 (Fig. S20f) mirrored Lanabecestat’s profile, with a dominant, well-defined low-energy basin, suggesting strong binding stabilization. This analysis reveals that phytochemicals, particularly Bacopaside 1, induce more structured and energetically favourable BACE1 conformations compared to Atabecestat and Verubecestat. The combination of stability (as seen in Bacopaside 1) and conformational diversity (exhibited by Bacopasaponin) suggests that *Bacopa monnieri* compounds may offer superior BACE1 inhibition. These findings highlight the potential of *Bacopa monnieri* phytochemicals, especially Bacopaside 1, as lead compounds for BACE1 inhibition. Their ability to induce stable yet flexible protein conformations could translate to more effective inhibition, potentially surpassing some established synthetic inhibitors. This study underscores the value of natural products in drug discovery and opens new avenues for developing Alzheimer’s disease therapeutics.

We further investigated the distance of the active site residue with important BACE1 sites after the binding event with respective ligands (see Supplementary Table S2). Notably, Bacopaside 1 demonstrated the closest approach to Asp228 in the critical flap region (R1) at 13.57Å, outperforming all synthetic inhibitors, including Verubecestat (13.06Å). This unprecedented interaction suggests a distinctive binding mode that could be pivotal for BACE1 inhibition. Furthermore, Bacopaside 1 exhibited a balanced interaction profile across all examined BACE1 regions, maintaining competitive distances to both catalytic residues (Asp32 and Asp228) throughout the protein structure. In the 10S loop (R2), Bacopaside 1 showed distances (15.21Å to Asp32, 13.67Å to Asp228) comparable to the best-performing synthetic inhibitors. Remarkably, in Insert A (R4), Bacopaside 1 consistently outperformed all synthetic compounds, with distances of 21.08Å and 20.98Å to Asp32 and Asp228, respectively, compared to Verubecestat’s 21.73Å and 22.44Å. This balanced yet strong interaction profile distinguishes Bacopaside 1 from both other phytochemicals and synthetic inhibitors. While Bacopasaponin D often showed the closest interactions (Fig. 11e), Bacopaside 1’s consistent performance across all regions suggests a unique mechanism of action (Fig. 11f). Its ability to maintain close interactions with both catalytic residues across BACE1’s structure indicates potential conformational modulation. These findings position Bacopaside 1 as a promising lead compound for BACE1 inhibition. Its distinctive interaction pattern, particularly the unprecedented flap region affinity and balanced profile across BACE1’s structure, suggests a novel mechanism of action that could overcome limitations of current synthetic inhibitors.

**Figure 11:**
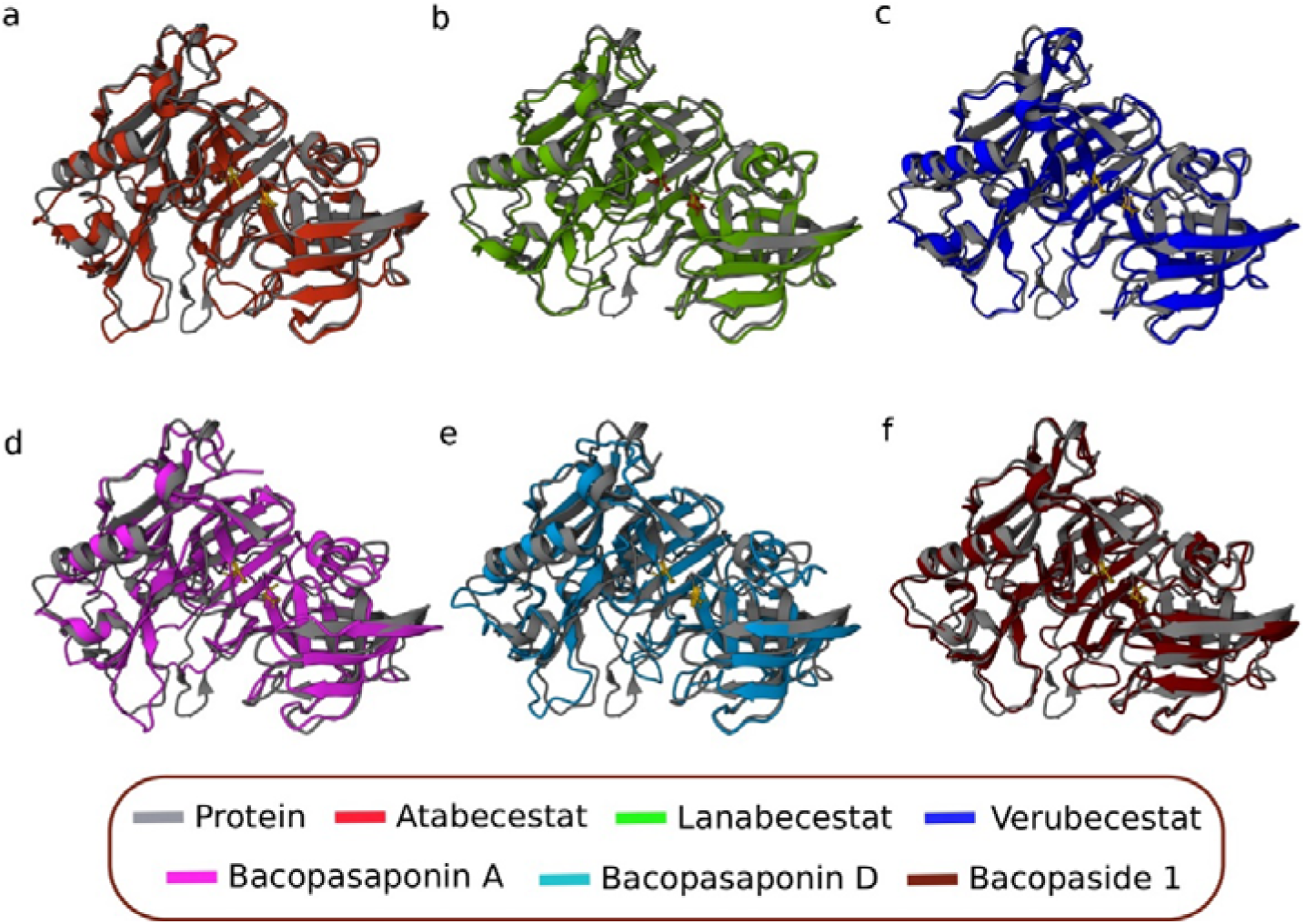
Structure of BACE1 simulated with respective ligands overlapped with apo form of BACE1 (grey). The panel demonstrates the structural changes in BACE1 when in contact with (a) Atabecestat (b) Lanabecestat (c) Verubecestat (d) Bacopasaponin A (e) Bacopasaponin D (f) Bacopaside 1

### SIMANA (SIMulation ANAlysis)

The comprehensive analyses we have conducted on the dynamic interactions between BACE1 and various ligands have culminated in the development of a powerful visualization platform called SIMANA (***SIMulation ANAlysis,*** https://simana.streamlit.app/). This web-based tool empowers researchers to easily explore and interpret molecular dynamics simulation data, providing a seamless interface for studying the structural and functional aspects of drug-target complexes. The journey that led to the creation of SIMANA began with our meticulous examination of clinically tested BACE1 inhibitors and the promising Bacopa monnieri phytochemicals. Through a multi-faceted approach involving several analyses, we uncovered compelling evidence that these natural compounds possess the potential to be more effective and specific BACE1 inhibitors compared to their synthetic counterparts. Recognizing the value of these insights, we were inspired to create a user-friendly platform that would enable researchers to visualize and interpret simulation-derived data with ease. Thus, SIMANA was created, a web-based tool that seamlessly integrates the visualization techniques, making them accessible to the broader scientific community. Through SIMANA, researchers can now easily generate high-quality, interactive visualizations of key simulation metrics, such as RMSD, RMSF, radius of gyration, SASA, dynamic cross-correlation matrices, and principal component analysis. The platform’s intuitive interface and customizable display options allow users to explore these parameters in depth, fostering a deeper understanding of the structural dynamics and conformational changes within protein-ligand complexes. While SIMANA primarily focuses on providing visualization capabilities, it also offers a suite of computational tools for additional analyses, such as Ramachandran plot generation, Lipinski’s rule calculations, and B-factor analysis. These complementary features enable researchers to gain a more comprehensive understanding of the physicochemical properties and structural characteristics of their systems of interest. By empowering researchers with the ability to visualize and interpret simulation data, SIMANA aims to accelerate the drug discovery process, facilitating the exploration of novel therapeutic candidates, including the promising *Bacopa monnieri* phytochemicals we have identified.

## Conclusion

The pursuit of effective BACE1 inhibitors remains a critical endeavour in the fight against Alzheimer’s disease, a condition that continues to pose significant challenges to global health. In this study, we explored the potential of natural phytochemicals derived from *Bacopa monnieri* as alternative therapeutic agents, juxtaposing their efficacy against established synthetic inhibitors. Our findings reveal that these natural compounds not only exhibit promising binding affinities but also demonstrate unique mechanisms of interaction with BACE1, suggesting a multifaceted approach to inhibition that could enhance therapeutic outcomes. The comparative analysis of the selected phytochemicals, including Bacopaside I, against known BACE1 inhibitors such as Atabecestat, Lanabecestat, and Verubecestat, underscores the potential of these natural compounds. The molecular docking studies indicated that Bacopaside I, in particular, adopts a more compact binding mode, which may correlate with a higher binding affinity. This observation is significant, as it suggests that natural compounds can achieve comparable, if not superior, efficacy to synthetic drugs, thereby warranting further investigation into their pharmacological properties. Moreover, Bacopaside I exhibits a remarkable binding affinity to BACE1, achieving a distance of 13.57 Å to Asp228 in the flap region, which is competitive with Verubecestat’s 13.06 Å. In the 10S loop, Bacopaside I maintains favourable distances of 15.21 Å to Asp32 and 13.67 Å to Asp228, comparable to the best synthetic inhibitors. Its performance in Insert A is particularly noteworthy, with distances of 21.08 Å and 20.98 Å to the catalytic residues, surpassing Verubecestat’s measurements. These consistent and close interactions across critical regions suggest that Bacopaside I may offer a unique mechanism of action that enhances its inhibitory potential. Overall, Bacopaside I positions itself as a promising alternative to Verubecestat, potentially providing superior therapeutic benefits in BACE1 inhibition. As the limitations of synthetic drugs become increasingly apparent, the resurgence of interest in phytochemicals offers a promising avenue for the development of novel therapeutic candidates. The bioactive compounds identified in this study not only provide a foundation for future research but also highlight the importance of traditional medicine in contemporary drug discovery. In conclusion, our investigation into the BACE1 inhibitory potential of *Bacopa monnieri* phytochemicals presents compelling evidence for their efficacy as viable alternatives to synthetic inhibitors. The promising results obtained from molecular docking studies, coupled with the innovative visualization capabilities of the SIMANA platform (https://simana.streamlit.app/), pave the way for further exploration of these natural compounds. As we continue to unravel the complexities of Alzheimer’s disease, the integration of natural products into therapeutic strategies may offer a beacon of hope in the quest for effective treatments. Future studies should focus on the in vivo efficacy and safety profiles of these phytochemicals, as well as their mechanisms of action, to fully elucidate their potential in combating neurodegenerative diseases.

## Methodology

### Ligand and Phytochemicals Selection

To investigate potential BACE1 inhibitors, we employed a comparative approach using both synthetic and natural compounds. Three clinically-studied BACE1 inhibitors—Atabecestat, Lanabecestat, and Verubecestat—were selected as reference ligands due to their established efficacy and progression to clinical trials for Alzheimer’s disease treatment. For exploring natural alternatives, we focused on eight specific bacosides from *Bacopa monnieri*, a plant traditionally used for cognitive enhancement: Bacopasaponin A, Bacopasaponin B, Bacopasaponin C, Bacopasaponin D, Bacopasaponin G, Bacopaside I, Bacopaside II, and Sarsasapogenin. These triterpenoid saponins, unique to *B. monnieri*, were chosen based on their prevalence in the plant and previous studies suggesting neuroprotective properties^29^.

#### Molecular Docking

The BACE1 active site, comprising the catalytic dyad (Asp32 and Asp228) and surrounding residues (e.g., Thr72, Ser35, Gly34, Arg235, and Tyr71), was analysed for ligand interactions. The structures of the inhibitors and selected phytochemicals were obtained from the PubChem database (https://pubchem.ncbi.nlm.nih.gov/)^30^ in .sdf format and converted to .pdb format using OpenBabel (https://openbabel.org/)^32^. The ligand structures were then subjected to energy minimization using the MM2 algorithm^33^, as implemented in the Chem3D software^34^. Molecular docking studies employed three platforms: AutoDock^35^, DockThor^36^, and CBDock^37^, enhancing result robustness. For receptor preparation, the structure of BACE1 (PDB ID: 2OHM) was obtained from the RCSB Protein Data Bank^38^. Preparation involved the removal of unwanted residues (small ions) and water molecules, followed by energy minimization using Chimera^39^. The minimization was conducted using the Steepest Descent Algorithm with 1000 steps and a step size of 0.1 Å, followed by 100 conjugate gradient steps with a step size of 0.1 Å to ensure optimal energetic conformation. The default AMBER ff14SB force field^40^ was applied to standard residues, and the AMI-BCC force field was used for other residues. Ligand-receptor interactions were visualized using LigPlot^41^. The docking result for each docking procedure are provided in the Supplementary File (see Supplementary Table S1). Redocking was performed by docking N∼3∼-benzylpyridine-2,3-diamine (inhibitor present in 2OHM PDB structure) with the BACE1 crystal structure to validate the docking protocol. Validation was confirmed by calculating the RMSD between the docked and crystal-bound N∼3∼-benzylpyridine-2,3-diamine molecules.

### Molecular Dynamic Simulation

Molecular dynamics (MD) simulations were performed to explore the dynamic interactions of known inhibitors and the top-ranked phytochemicals with BACE1. All simulations were executed in triplicate using GROMACS version 2018.1^42,43,44^ for a duration of 150 ns. The protein-ligand complexes were prepared with the CHARMM36 force field, and ligand topologies were generated using CGenFF^45^. The simulation system was solvated in a 5 nm³ cubic box with the TIP3P water model, and charge neutrality was achieved by adding Na^+^ and Cl^-^ ions. Energy minimization was carried out using the steepest descent algorithm for a maximum of 5000 steps. The system was then equilibrated through two consecutive 1 ns step: first under an NVT ensemble at 300K using a V-rescale thermostat, followed by an NPT ensemble at 1 bar pressure using a Berendsen barostat. The production MD phase ran for 150 ns using the leap-frog algorithm with a 2-fs time step. Van der Waals interactions were treated with a 1.2 nm cutoff, while long-range electrostatic interactions were computed using Particle Mesh Ewald (PME). The Nosé-Hoover thermostat and the Parrinello-Rahman barostat were applied to maintain temperature and pressure stability throughout the simulation. Coordinates were saved every 20 ps for subsequent analysis. Post simulation analyses were carried out to assess the stability, flexibility, and conformational behaviour of the protein-ligand complexes.

### Molecular Mechanics Possoin-Boltzmann Surface Area (MM-PBSA)

The calculation of binding free energy (ΔG_bind_) between a protein and its ligand is crucial for understanding the molecular interactions within protein-ligand complexes. In this study, we employed the Molecular Mechanics/Poisson-Boltzmann Surface Area (MM/PBSA) approach, a widely accepted method for estimating binding free energy. In MM/PBSA, the binding free energy (ΔG_bind_) is derived as the difference between the free energy of the protein-ligand complex (ΔG_complex_) and the sum of the free energies of the isolated protein (ΔG_protein_) and ligand (ΔG_ligand_ ). This relationship is mathematically expressed as:

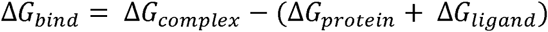

Here, ΔG_complex_ represents the free energy of the protein-ligand complex, while ΔG_protein_ and ΔG_ligand_ correspond to the free energies of the isolated protein and ligand, respectively.

This method provides a detailed energetic perspective on the stability and binding affinity of the complexes.

### Dynamic Cross Correlation Matrix (DCCM) Analysis

Dynamic cross-correlation analysis (DCCM) was conducted to explore the associations between protein residues. A structural ensemble was generated from the MD simulation trajectories of the protein in complex with the top-ranked ligands. A matrix representing these correlations was generated, with elements visualized as a Dynamic Cross-Correlation Matrix (DCCM). To enhance the robustness of the analysis, we utilized three independent MD simulation trajectories (triplicates), averaged the results, and calculated the final DCCM.

### Deformability Analysis

To assess the deformability and structural stability of BACE1 in the presence of the inhibitors and phytochemicals, Normal Mode Analysis (NMA) was performed using the iMOD server^46^. The equilibrated structures from the MD simulations were used as input. NMA was carried out by constructing an elastic network model of the protein, where each Cα atom is treated as a node, and the interatomic interactions are represented as springs. The server computed the collective motions of the protein along the lowest-frequency normal modes. Deformability analysis was conducted to calculate the deformation energy for each residue, indicating the ability of the residues to undergo conformational changes. Regions with high deformability correspond to areas with greater flexibility, while those with lower deformability represent rigid regions.

### Principal Component Analysis (PCA)

Principal Component Analysis (PCA) was employed to elucidate the structural dynamics of the protein-ligand complex by analysing the contribution of specific residues to each principal component (PC). To achieve this, we computed the covariance matrix of the system, capturing the relationships between the positional fluctuations of individual residues. Eigen decomposition of the covariance matrix was then performed, yielding eigenvalues and corresponding eigenvectors. The eigenvalues represent the variance explained by each PC, while the eigenvectors indicate the contribution of individual residues to these components. We calculated the contribution of each residue to the PCs, allowing us to pinpoint which regions of the protein contribute most significantly to the observed structural variance. This approach enabled a residue-level interpretation of the principal components, offering deeper insights into the regions of the protein that are most dynamically involved in ligand binding.

## Notes

### Competing Interest Statement

The authors have declared no competing interest.

