## Supplementary file for "*Bacopa monnieri* phytochemicals as promising BACE1 inhibitors for Alzheimer’s Disease Therapy"

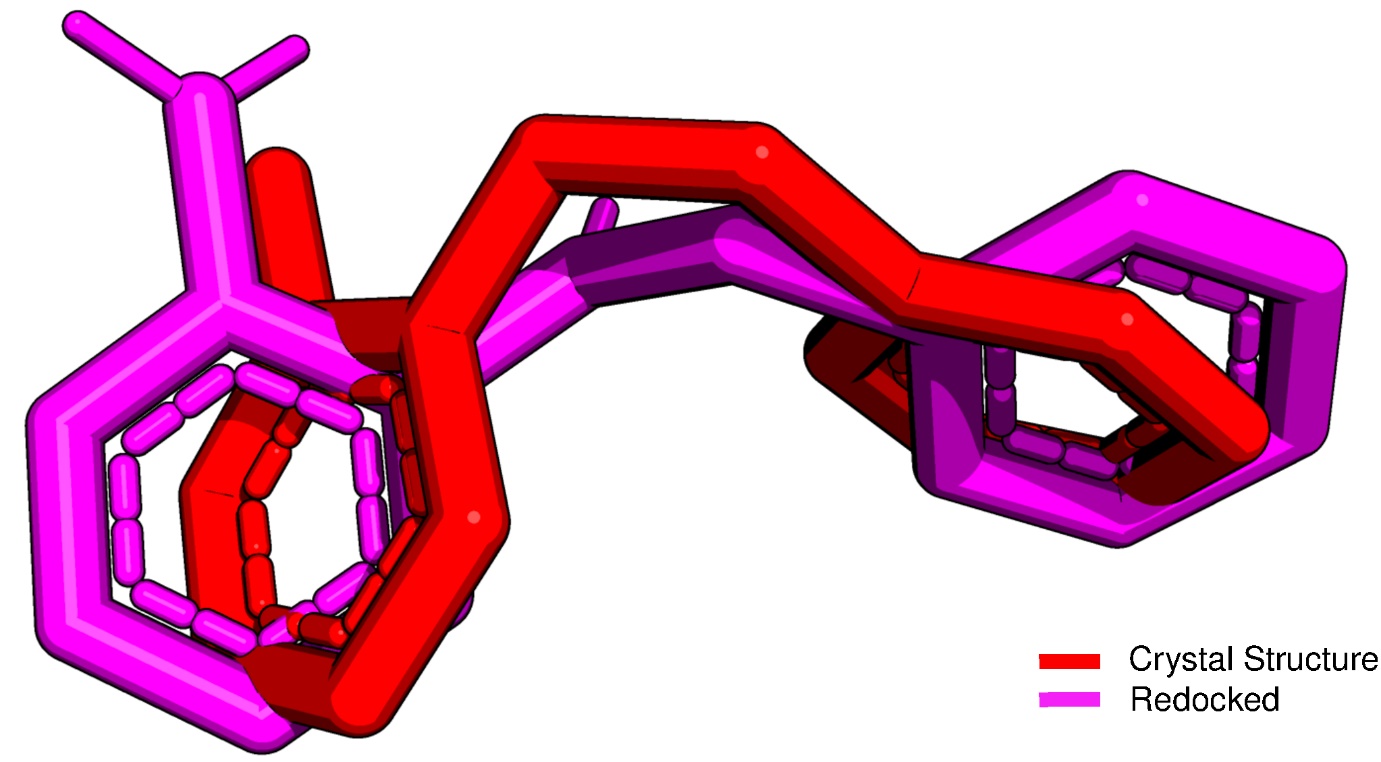

**Figure S1**: Crystal (red) and Redocked structure (magenta) of N~3~-benzylpyridine-2,3-diamine with an average RMSD of 1.1 Å.

**Table S1**: Molecular Docking of Ligands and Phytochemicals using AutoDock Vina, DockThor and CBDock

| Ligands | AutoDock Vina (kcal/mol) | DockThor (kcal/mol) | CBDock (kcal/mol) |
| --- | --- | --- | --- |
| Atabecestat | -7.5 | -7.7 | -9.3 |
| Lanabecestat | -8.8 | -8.962 | -8.7 |
| Verubecestat | -7.4 | -8.035 | -8.6 |
| Bacopasaponin A | -8.6 | -8.994 | -9.2 |
| Bacopasaponin B | -8.2 | -8.722 | -8.2 |
| Bacopasaponin C | -8.1 | -8.761 | -8.4 |
| Bacopasaponin D | -8.6 | -8.841 | -8.6 |
| Bacopasaponin G | -8.2 | -8.593 | -8.4 |
| Bacopaside 1 | -8.6 | -9.24 | -9.3 |
| Bacopaside 2 | -8.4 | -8.753 | -8.5 |
| Sarsasapogenin | -8.0 | -8.255 | -8.2 |

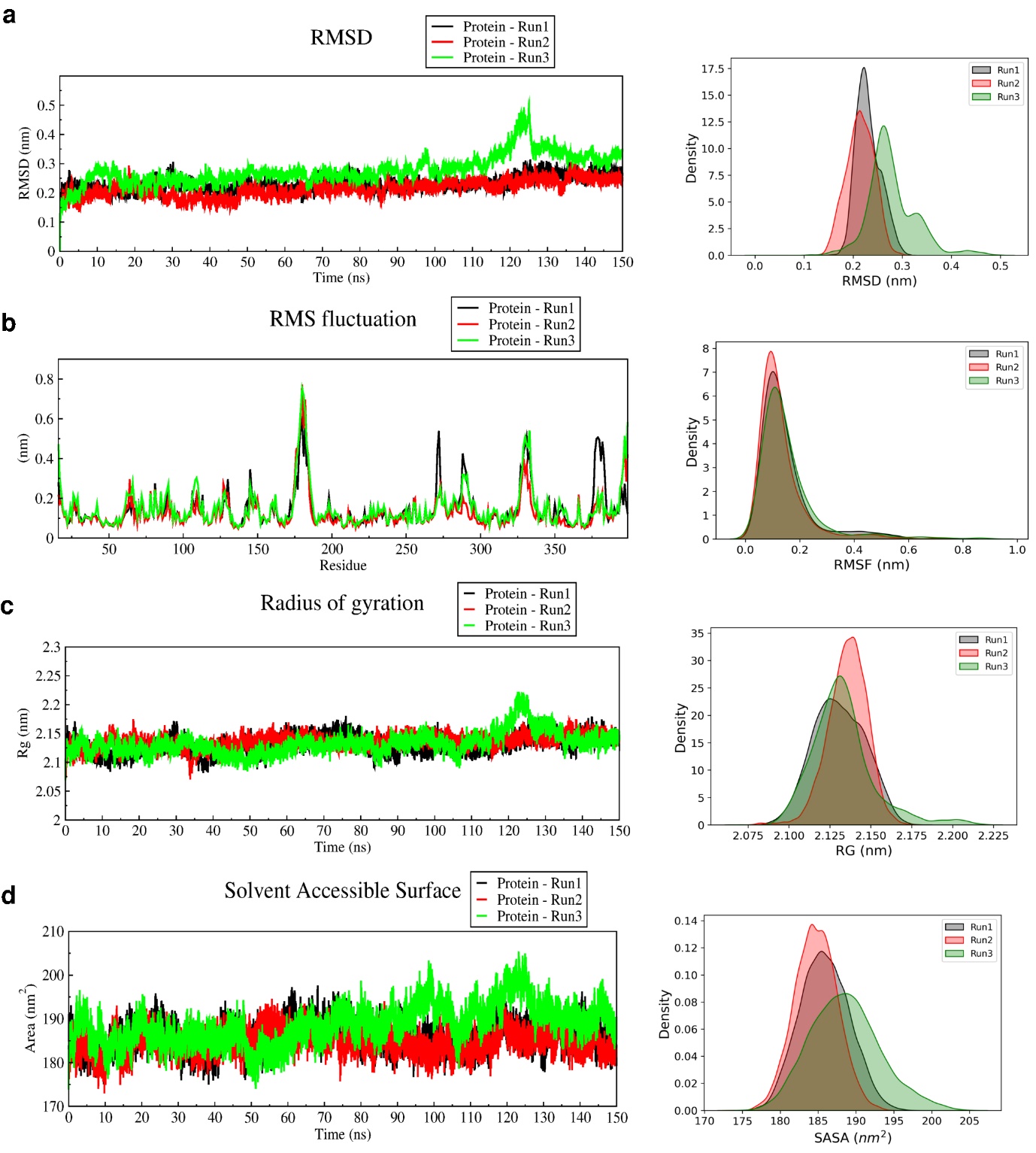

**Figure S2**: Triplicate MD simulation of Protein in apo form (no ligands) for 150 ns. (a) RMSD curve (left) and RMSD distribution (right) for three runs (b) RMSF curve (left) and RMSF distribution (right) for three runs (c) RG curve (left) and RG distribution (right) for three runs (a) SASA curve (left) and SASA distribution (right) for three runs

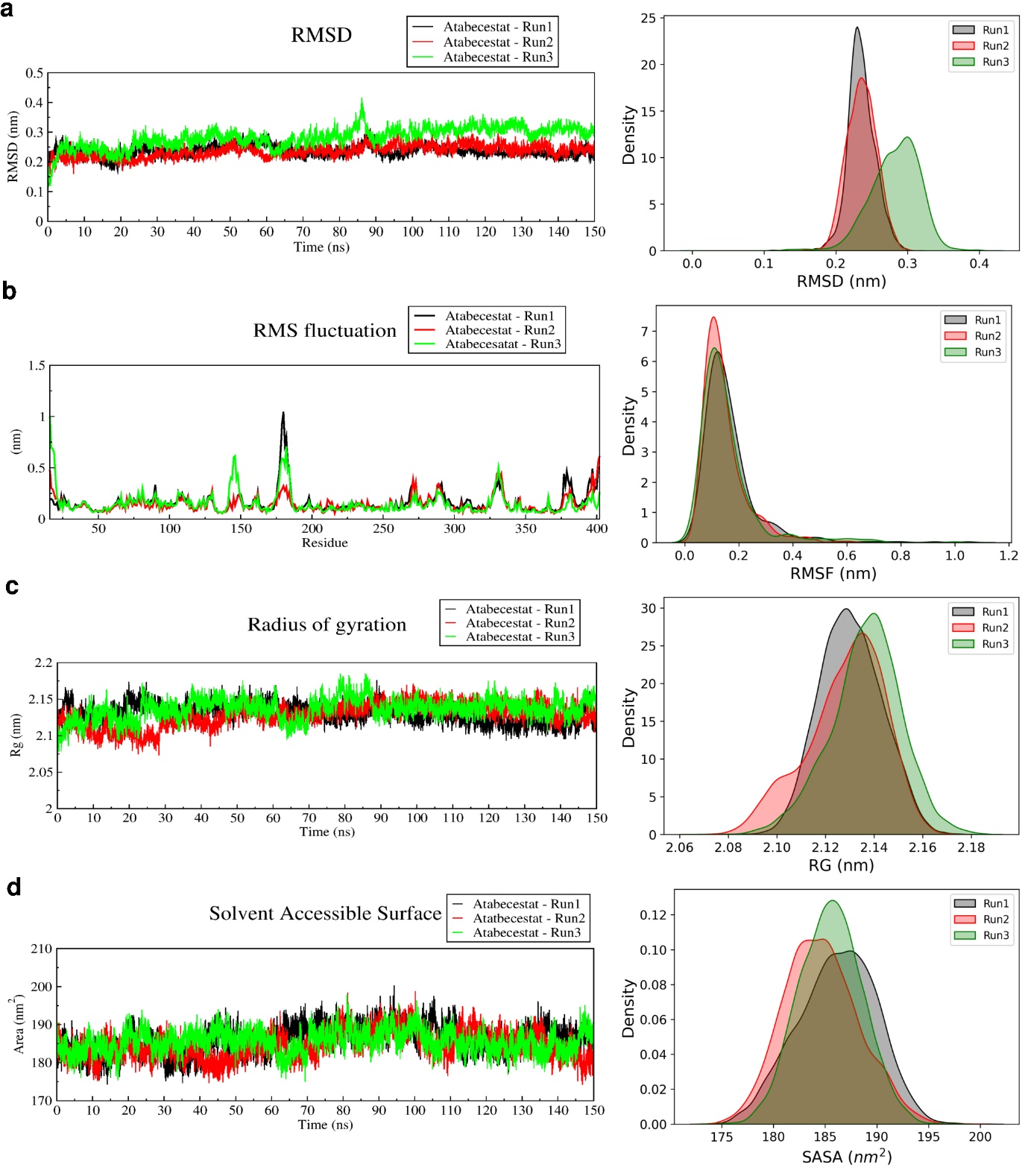

**Figure S3**: Triplicate MD simulation of Protein in apo form (no ligands) for 150 ns. (a) RMSD curve (left) and RMSD distribution (right) for three runs (b) RMSF curve (left) and RMSF distribution (right) for three runs (c) RG curve (left) and RG distribution (right) for three runs (a) SASA curve (left) and SASA distribution (right) for three runs

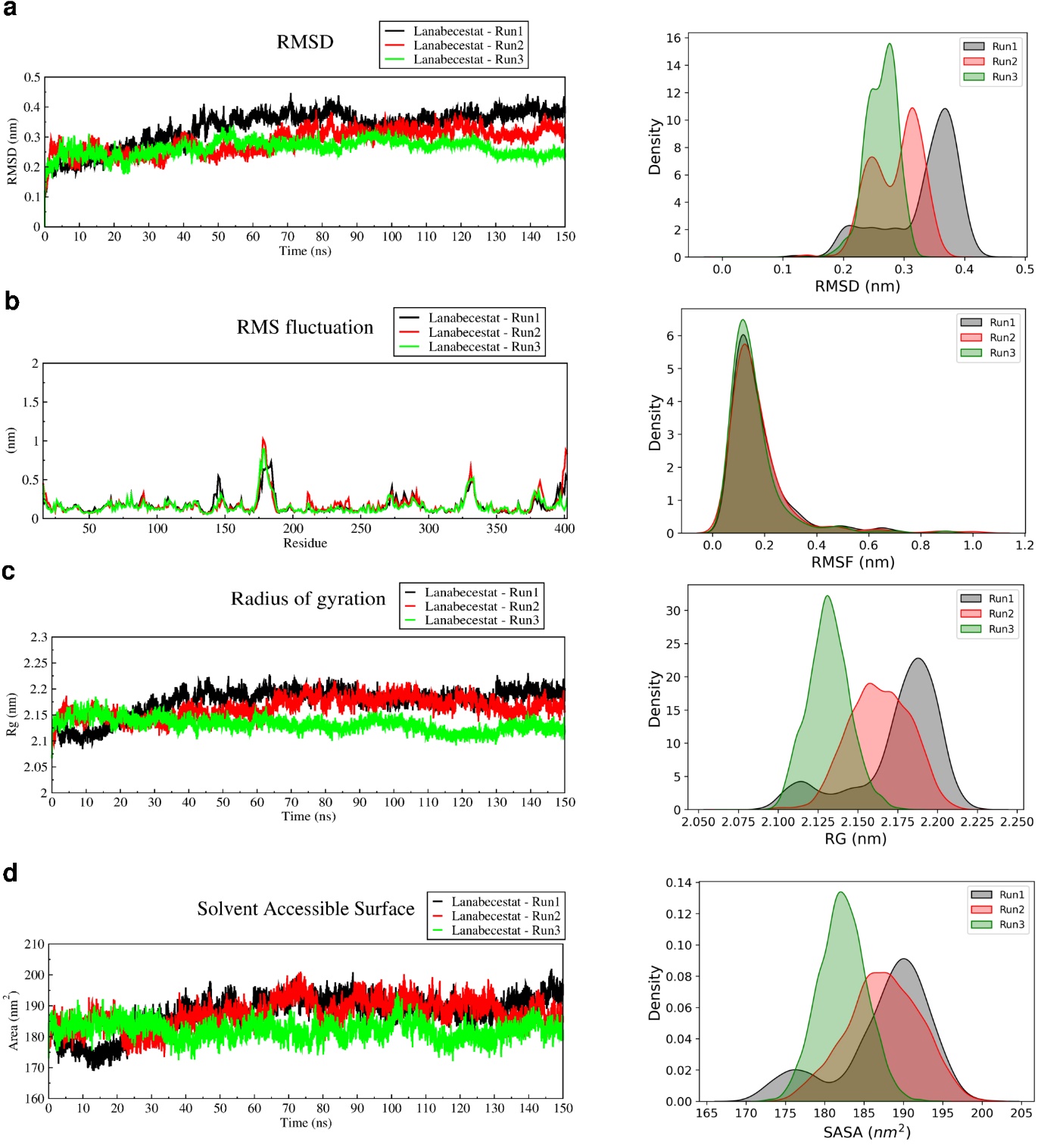

**Figure S4**: Triplicate MD simulation of Protein in apo form (no ligands) for 150 ns. (a) RMSD curve (left) and RMSD distribution (right) for three runs (b) RMSF curve (left) and RMSF distribution (right) for three runs (c) RG curve (left) and RG distribution (right) for three runs (a) SASA curve (left) and SASA distribution (right) for three runs

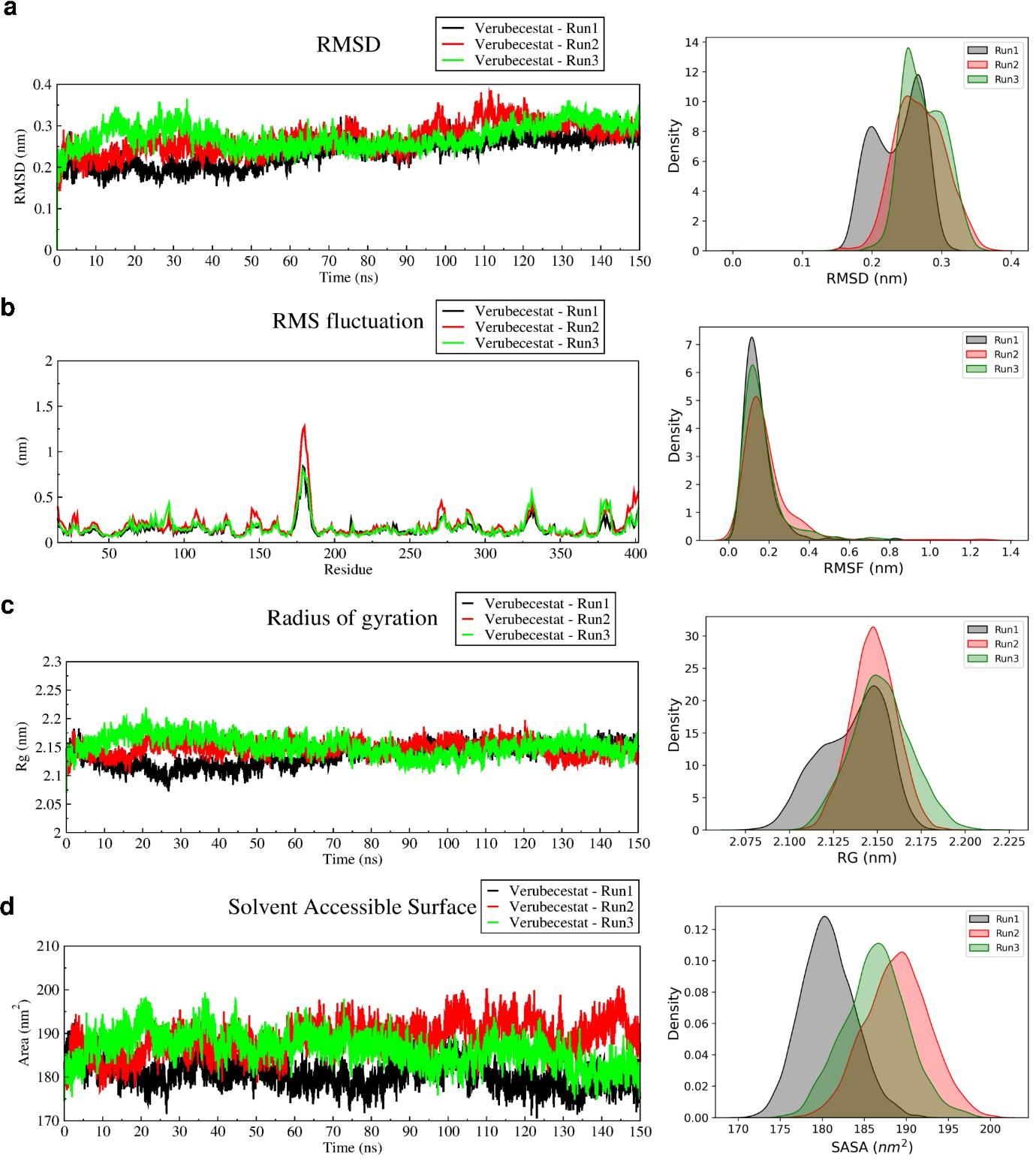

**Figure S5**: Triplicate MD simulation of Protein in apo form (no ligands) for 150 ns. (a) RMSD curve (left) and RMSD distribution (right) for three runs (b) RMSF curve (left) and RMSF distribution (right) for three runs (c) RG curve (left) and RG distribution (right) for three runs (a) SASA curve (left) and SASA distribution (right) for three runs

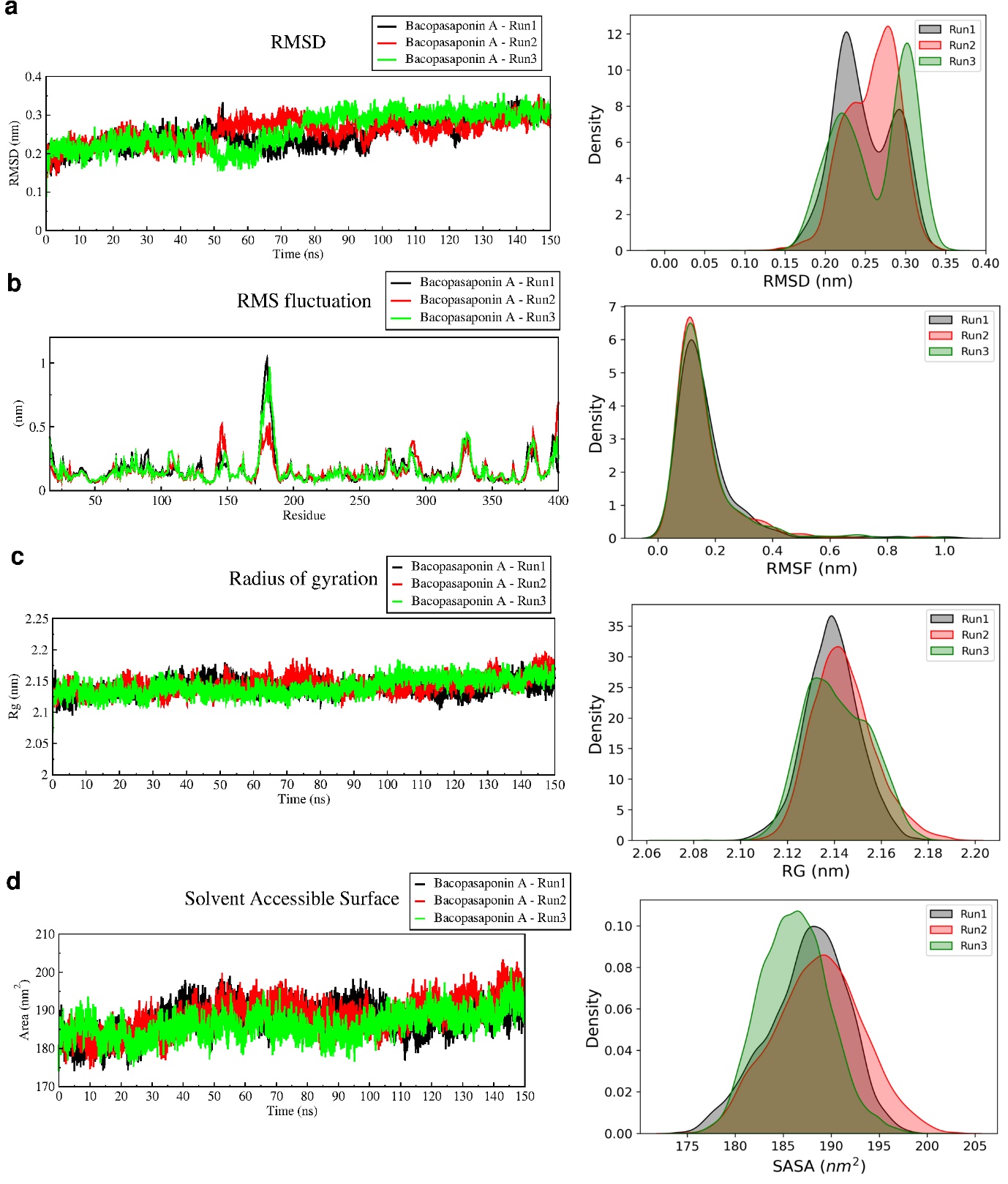

**Figure S6**: Triplicate MD simulation of Protein in apo form (no ligands) for 150 ns. (a) RMSD curve (left) and RMSD distribution (right) for three runs (b) RMSF curve (left) and RMSF distribution (right) for three runs (c) RG curve (left) and RG distribution (right) for three runs (a) SASA curve (left) and SASA distribution (right) for three runs

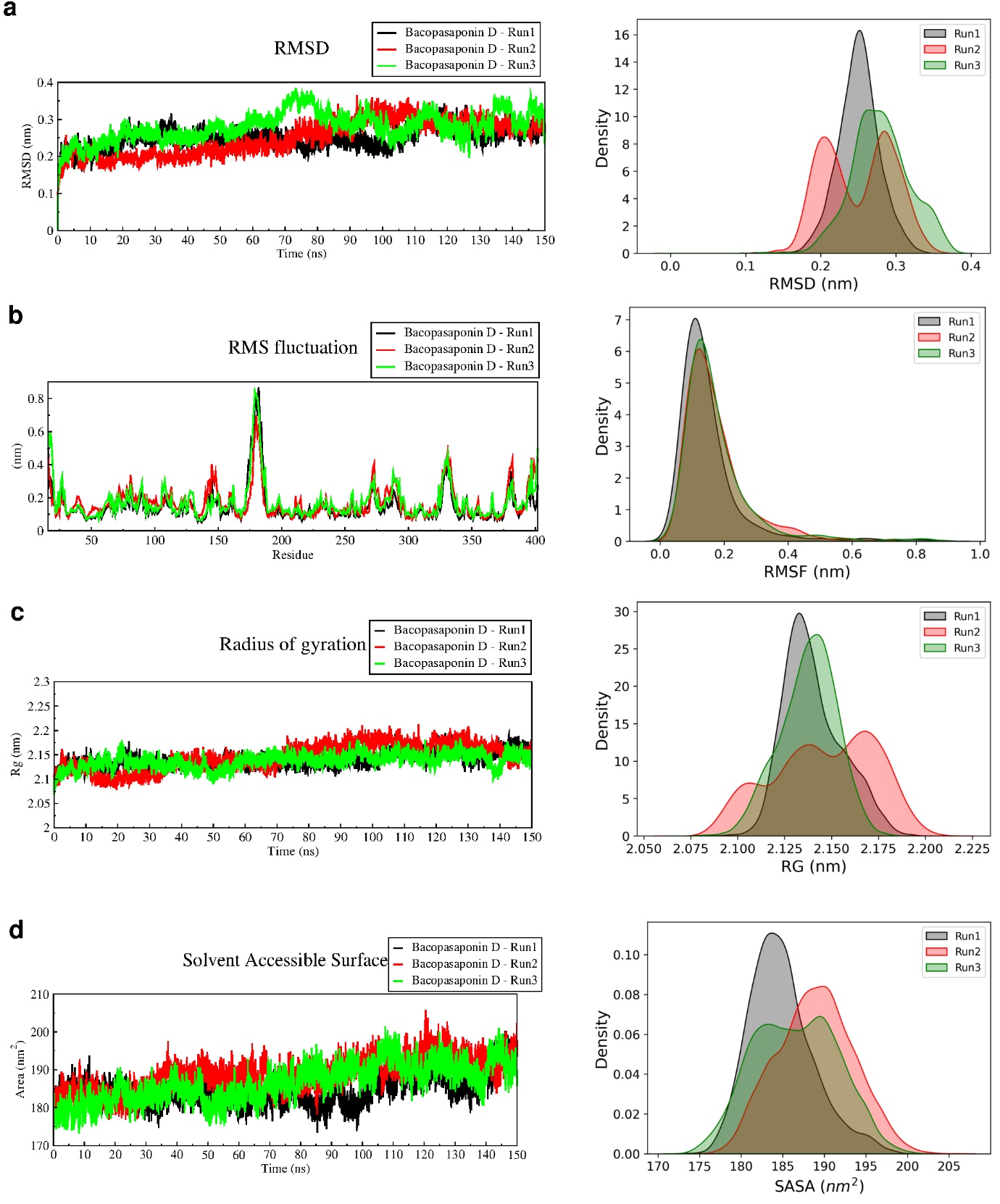

**Figure S7**: Triplicate MD simulation of Protein in apo form (no ligands) for 150 ns. (a) RMSD curve (left) and RMSD distribution (right) for three runs (b) RMSF curve (left) and RMSF distribution (right) for three runs (c) RG curve (left) and RG distribution (right) for three runs (a) SASA curve (left) and SASA distribution (right) for three runs

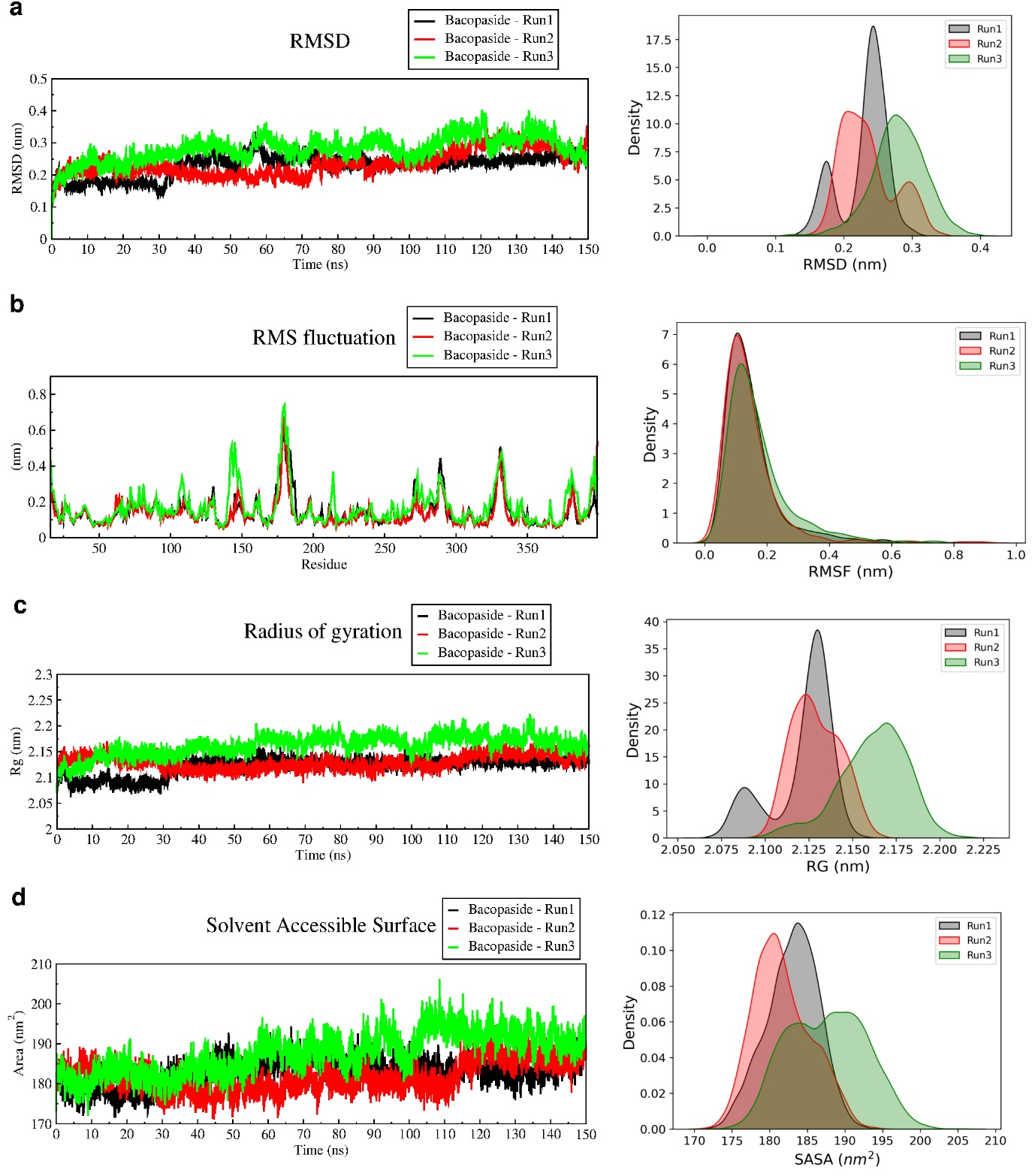

**Figure S8**: Triplicate MD simulation of Protein in apo form (no ligands) for 150 ns. (a) RMSD curve (left) and RMSD distribution (right) for three runs (b) RMSF curve (left) and RMSF distribution (right) for three runs (c) RG curve (left) and RG distribution (right) for three runs (a) SASA curve (left) and SASA distribution (right) for three runs

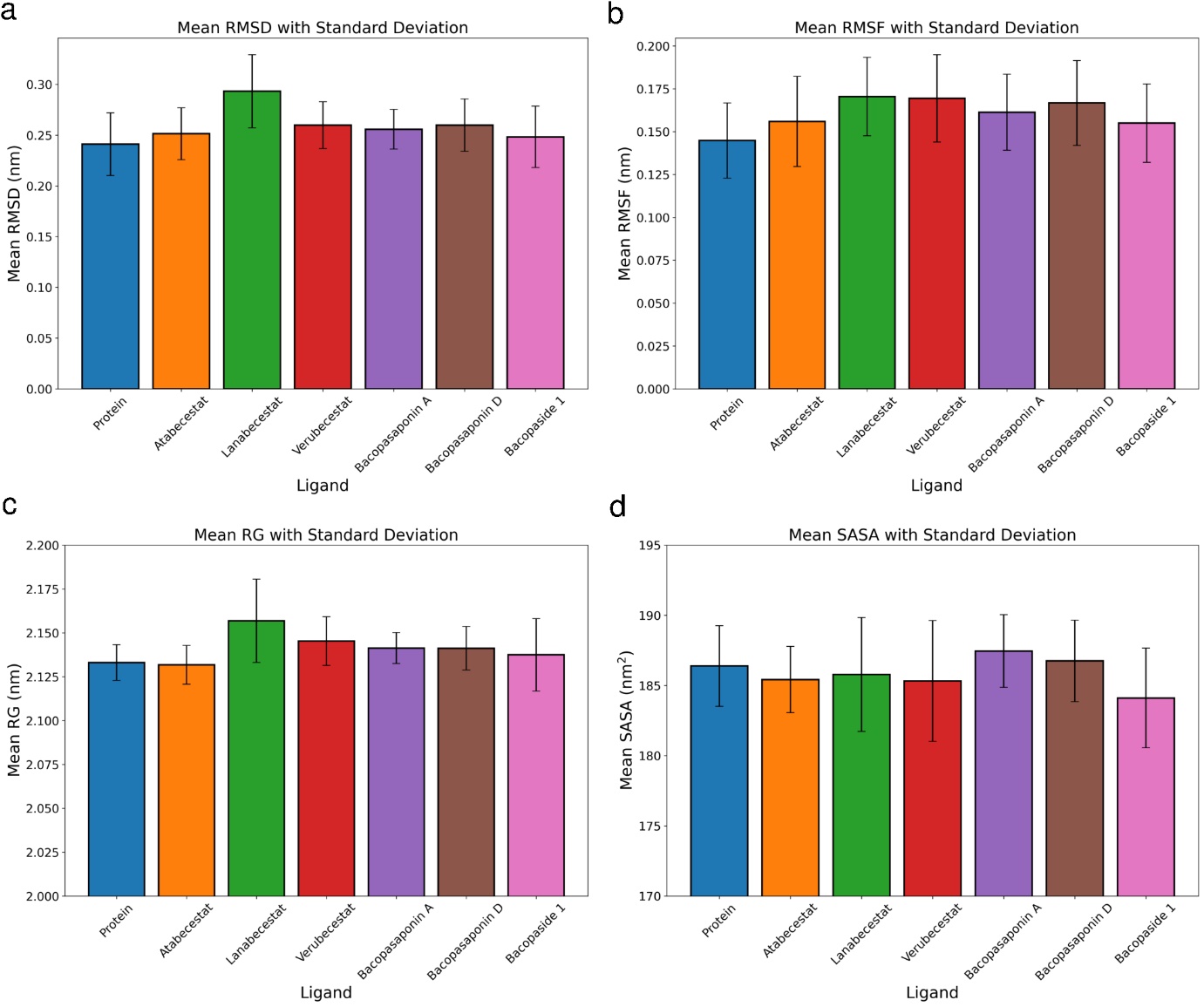

**Figure S9**: 150 ns Molecular Dynamic Simulation Statistical Analysis for (a) RMSD (b) RMSF (c) RG and (d) SASA

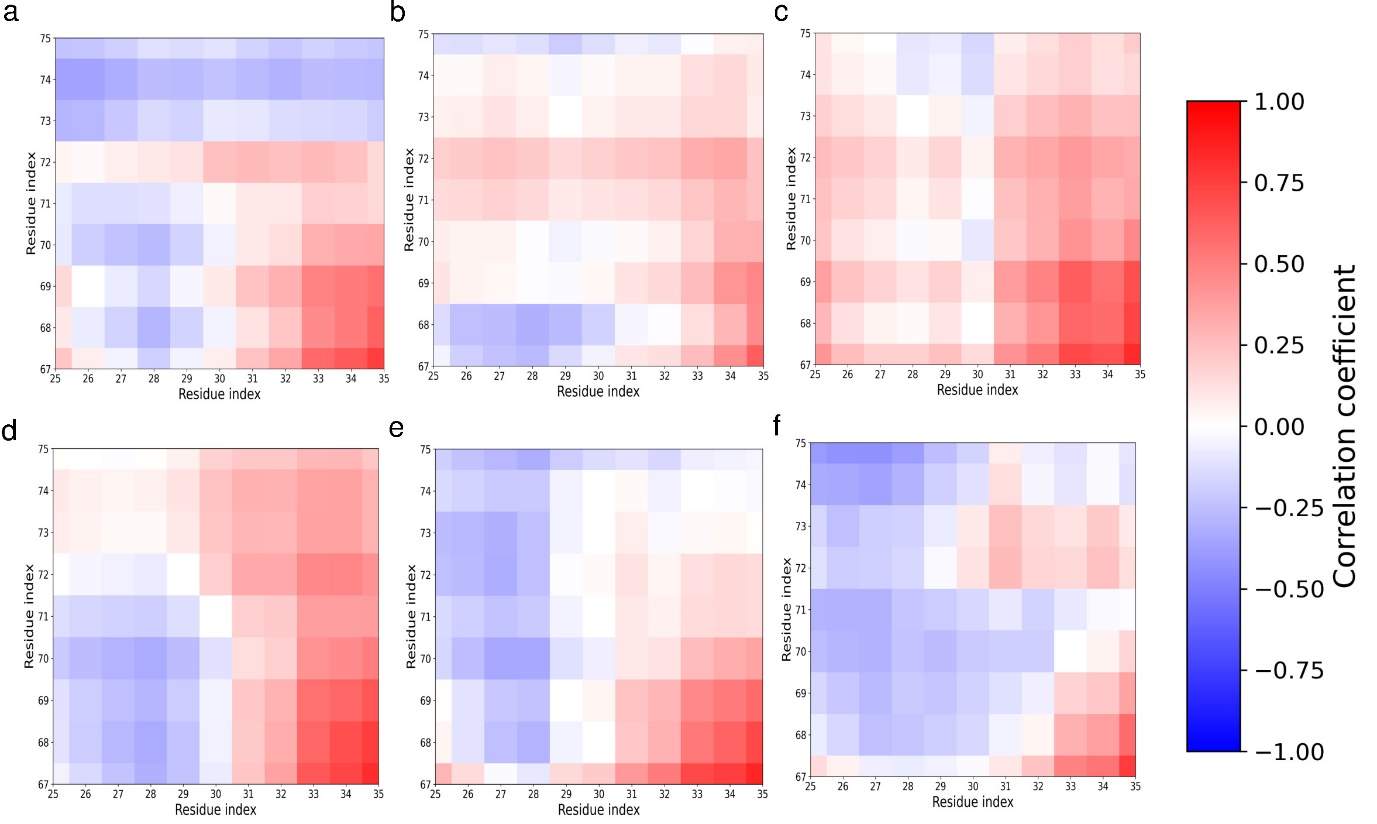

**Figure S10**: Dynamic Cross Correlation of BACE1 active site 1 (residue 25-35, including Asp32) and flap region (residues 67-75) with (a) Atabecestat (b) Lanabecestat (c) Verubecestat (d) Bacopasaponin A (e) Bacopasaponin D (f) Bacopaside 1

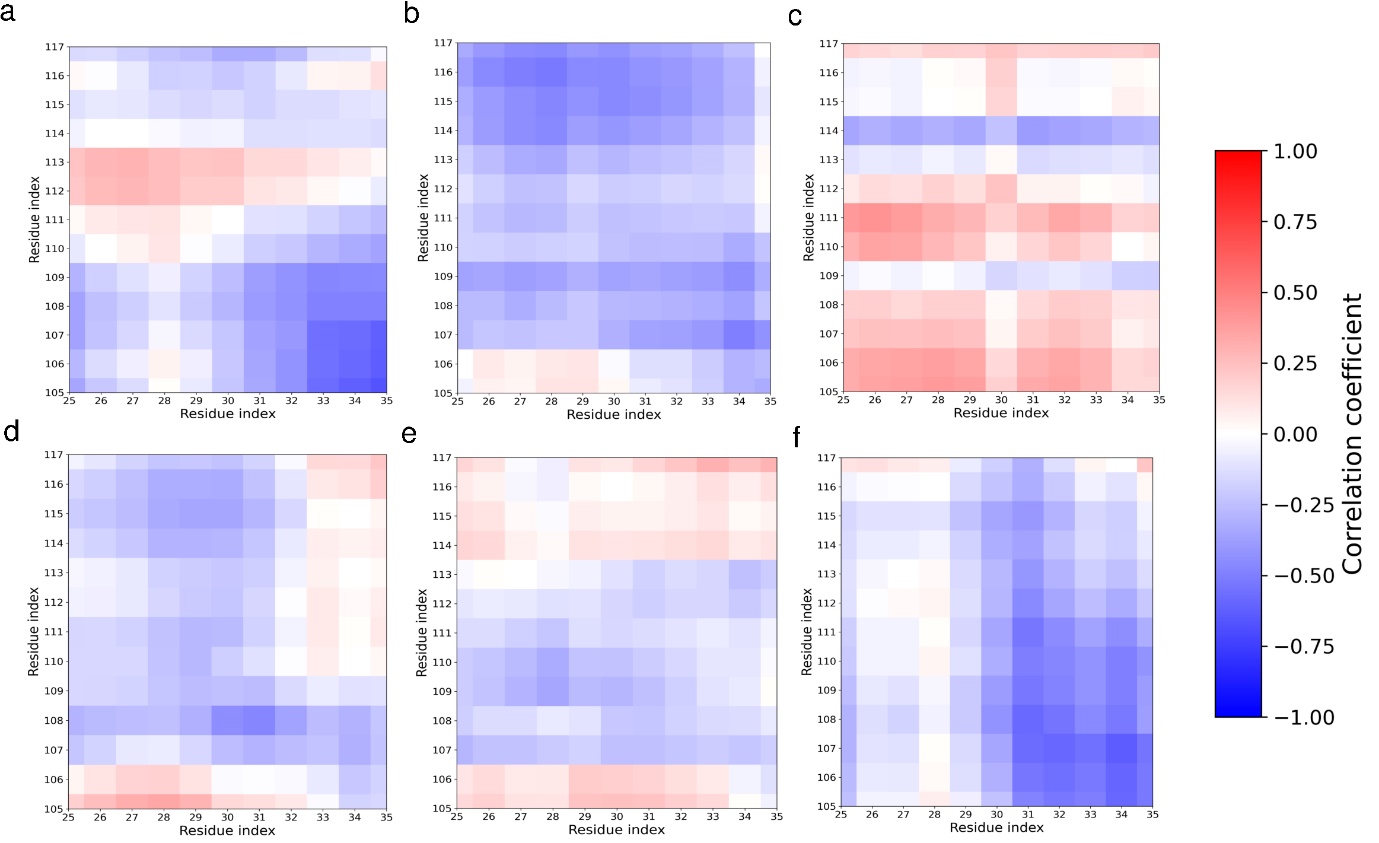

**Figure S11**: Dynamic Cross Correlation of BACE1 active site 1 (residue 25-35, including Asp32) and 113S loop region (residues 105-117) with (a) Atabecestat (b) Lanabecestat (c) Verubecestat (d) Bacopasaponin A (e) Bacopasaponin D (f) Bacopaside 1

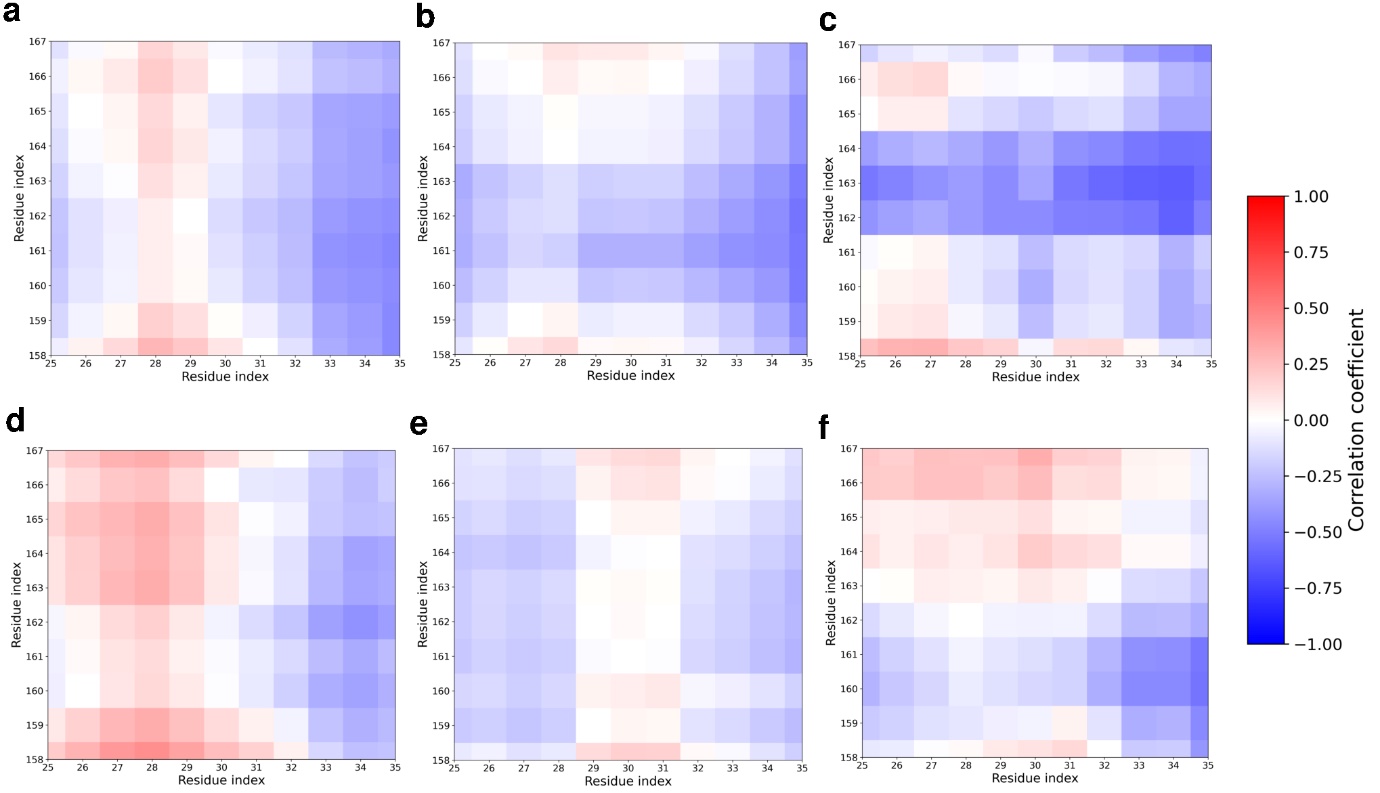

**Figure S12**: Dynamic Cross Correlation of BACE1 active site 1 (residue 25-35, including Asp32) and Insert A (residues 158-167) with (a) Atabecestat (b) Lanabecestat (c) Verubecestat (d) Bacopasaponin A (e) Bacopasaponin D (f) Bacopaside 1

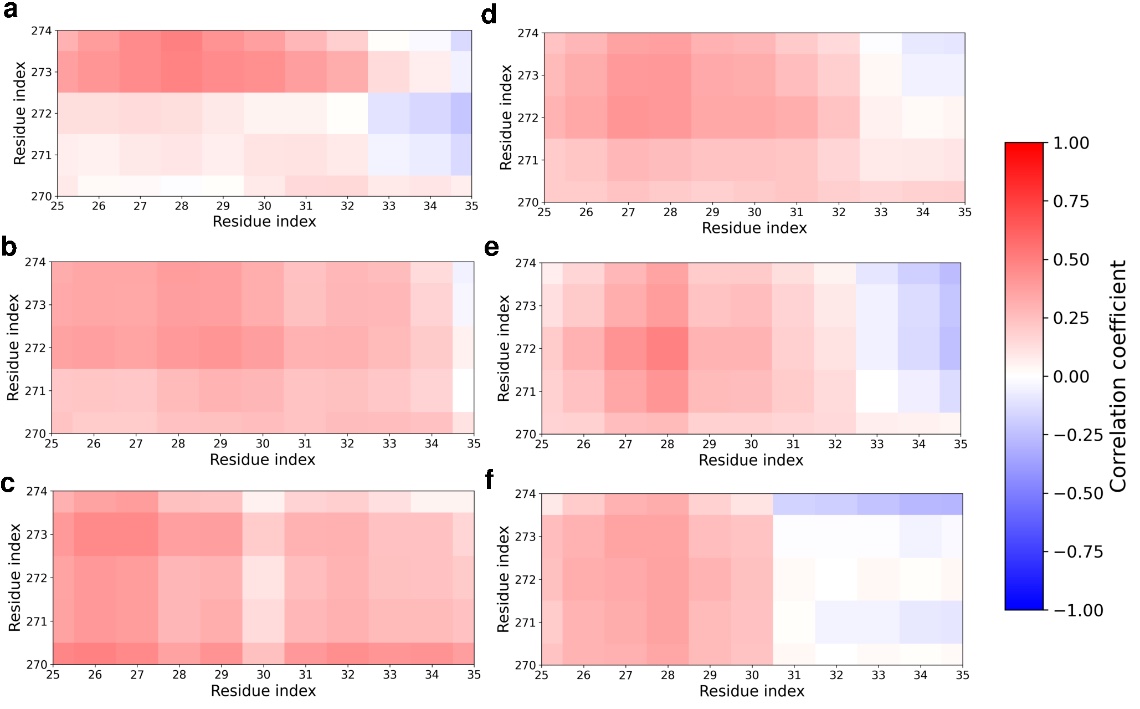

**Figure S13**: Dynamic Cross Correlation of BACE1 active site 1 (residue 25-35, including Asp32) and Insert D (residues 270-274) with (a) Atabecestat (b) Lanabecestat (c) Verubecestat (d) Bacopasaponin A (e) Bacopasaponin D (f) Bacopaside 1

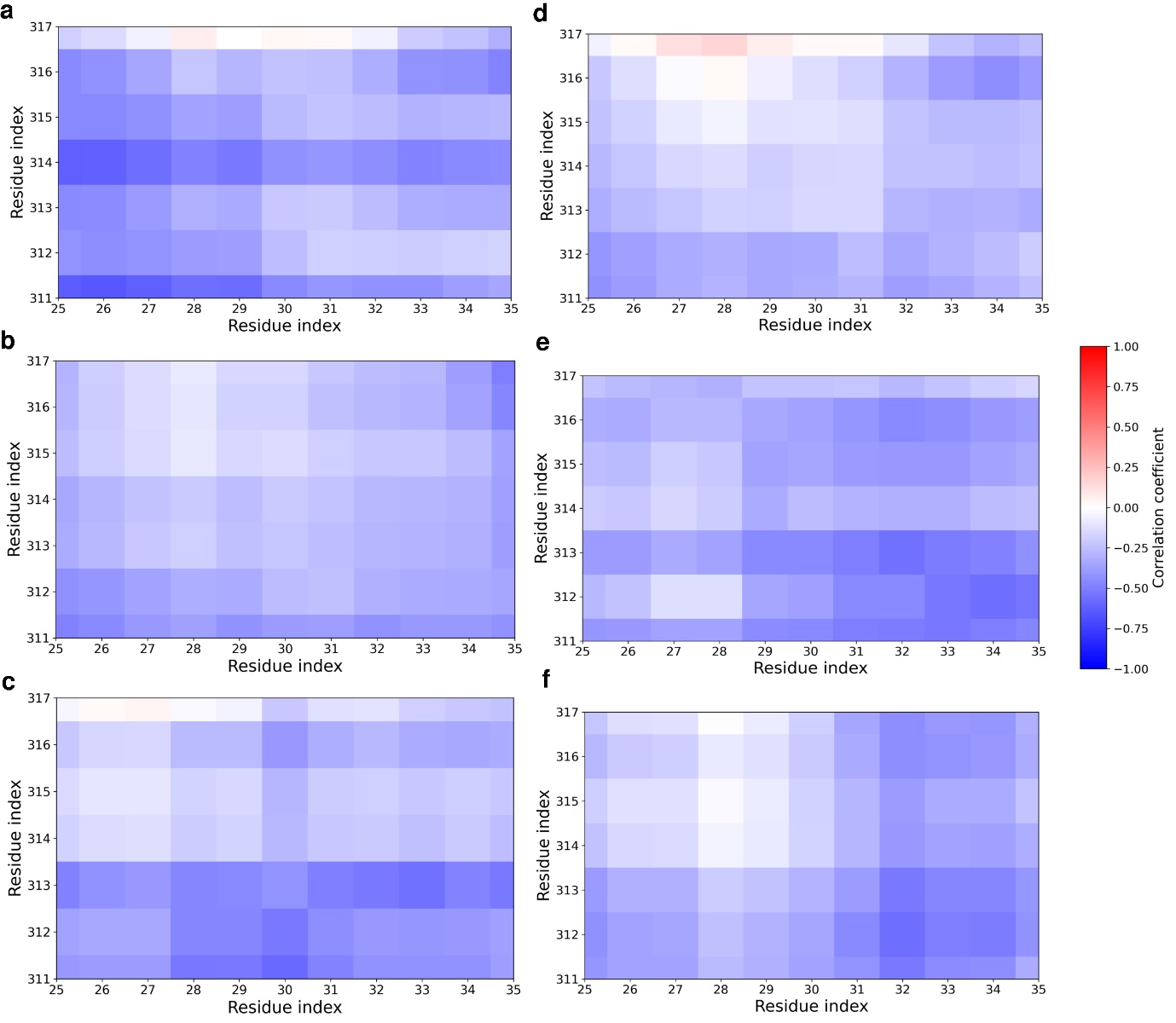

**Figure S14**: Dynamic Cross Correlation of BACE1 active site 1 (residue 25-35, including Asp32) and Insert F (residues 311-317) with (a) Atabecestat (b) Lanabecestat (c) Verubecestat (d) Bacopasaponin A (e) Bacopasaponin D (f) Bacopaside 1

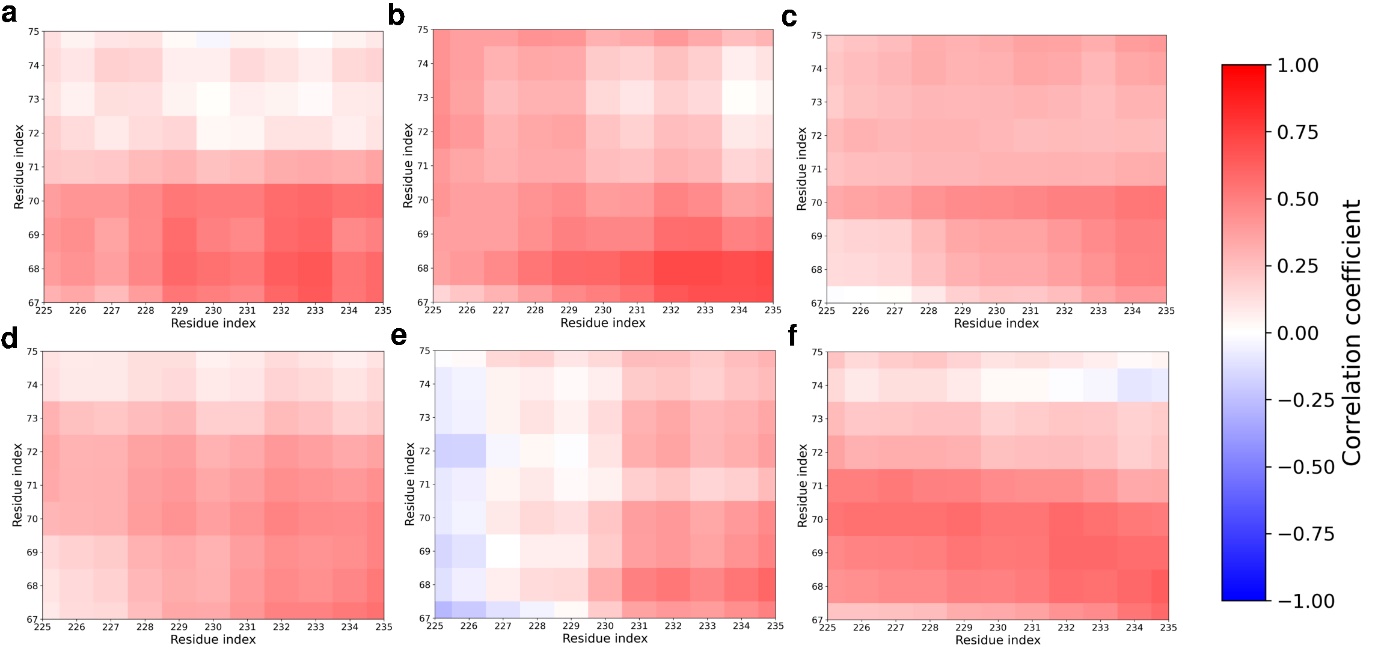

**Figure S15**: Dynamic Cross Correlation of BACE1 active site 2 (residue 225-235, including Asp228) and flap region (residues 67-75) with (a) Atabecestat (b) Lanabecestat (c) Verubecestat (d) Bacopasaponin A (e) Bacopasaponin D (f) Bacopaside 1

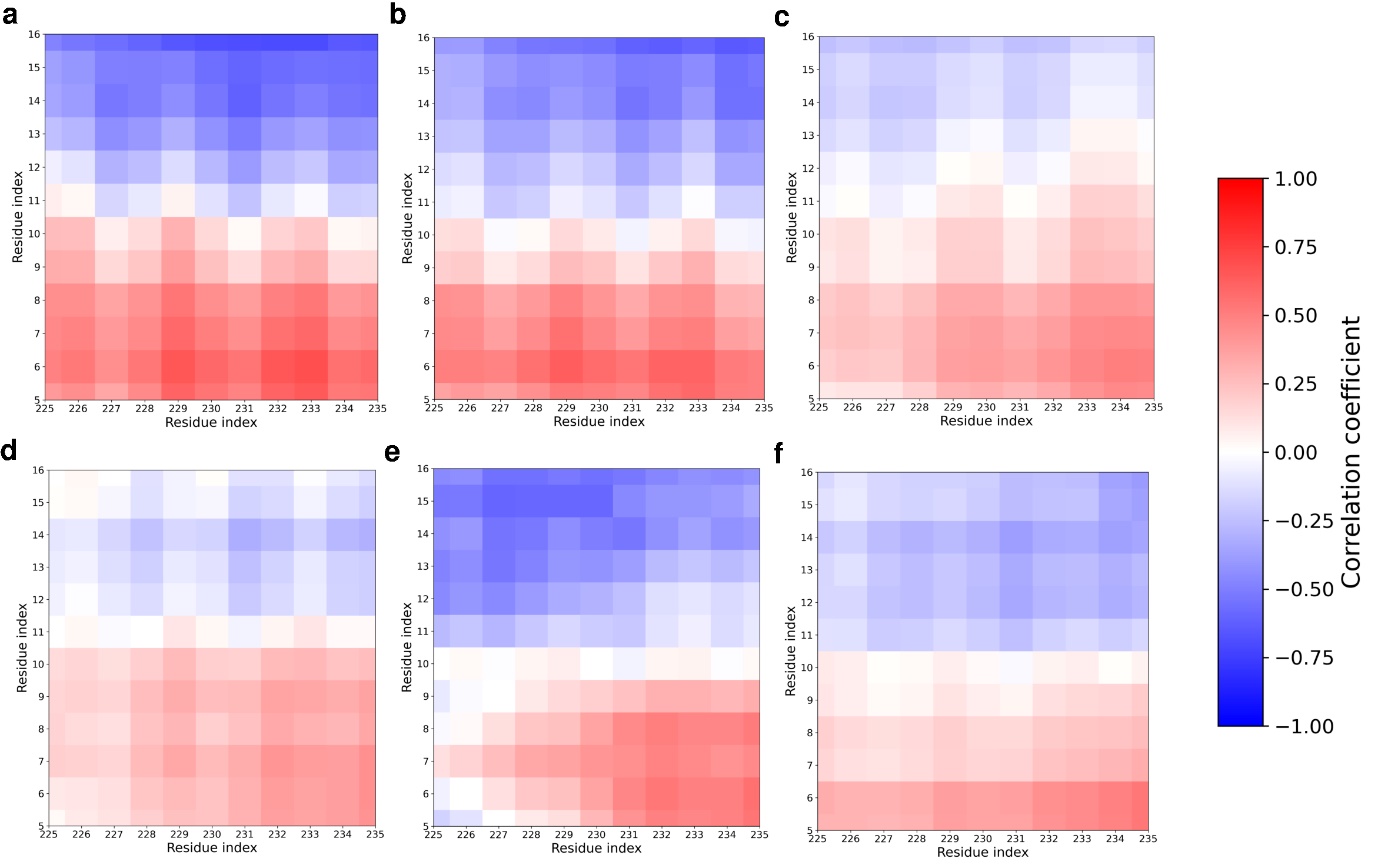

Figure 8: Dynamic Cross Correlation of BACE1 active site 2 (residue 225-235, including Asp32) and 10S loop region (residues 5-16) with (a) Atabecestat (b) Lanabecestat (c) Verubecestat (d) Bacopasaponin A (e) Bacopasaponin D (f) Bacopaside 1

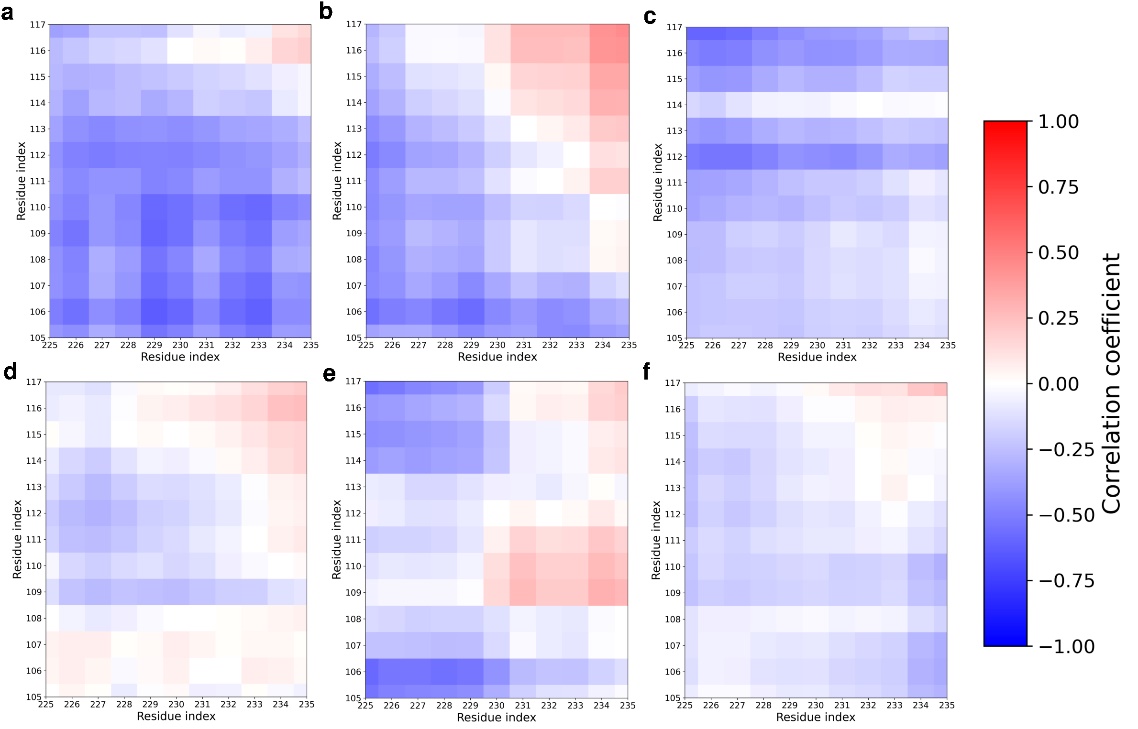

**Figure S16**: Dynamic Cross Correlation of BACE1 active site 2 (residue 225-235, including Asp228) and 113S loop (residues 105-117) with (a) Atabecestat (b) Lanabecestat (c) Verubecestat (d) Bacopasaponin A (e) Bacopasaponin D (f) Bacopaside 1

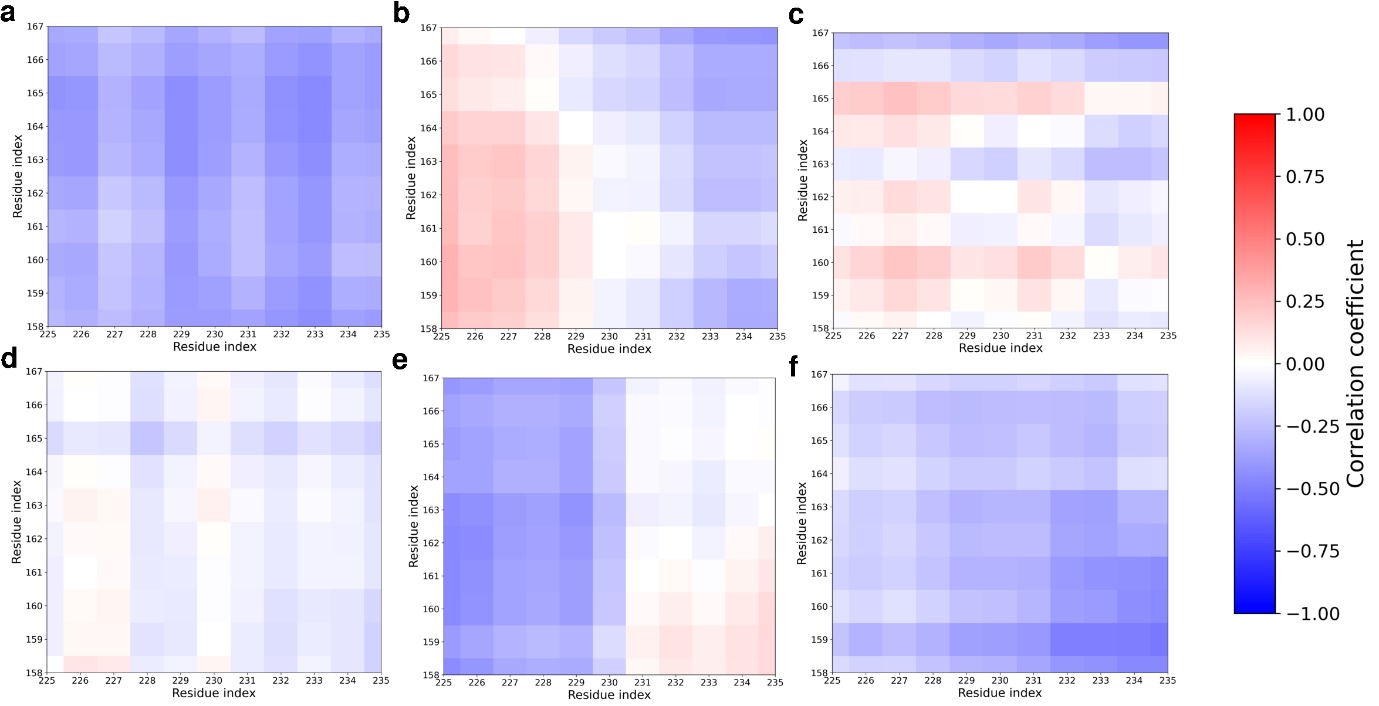

**Figure S17**: Dynamic Cross Correlation of BACE1 active site 2 (residue 225-235, including Asp228) and Insert A (residues 158-167) with (a) Atabecestat (b) Lanabecestat (c) Verubecestat (d) Bacopasaponin A (e) Bacopasaponin D (f) Bacopaside 1

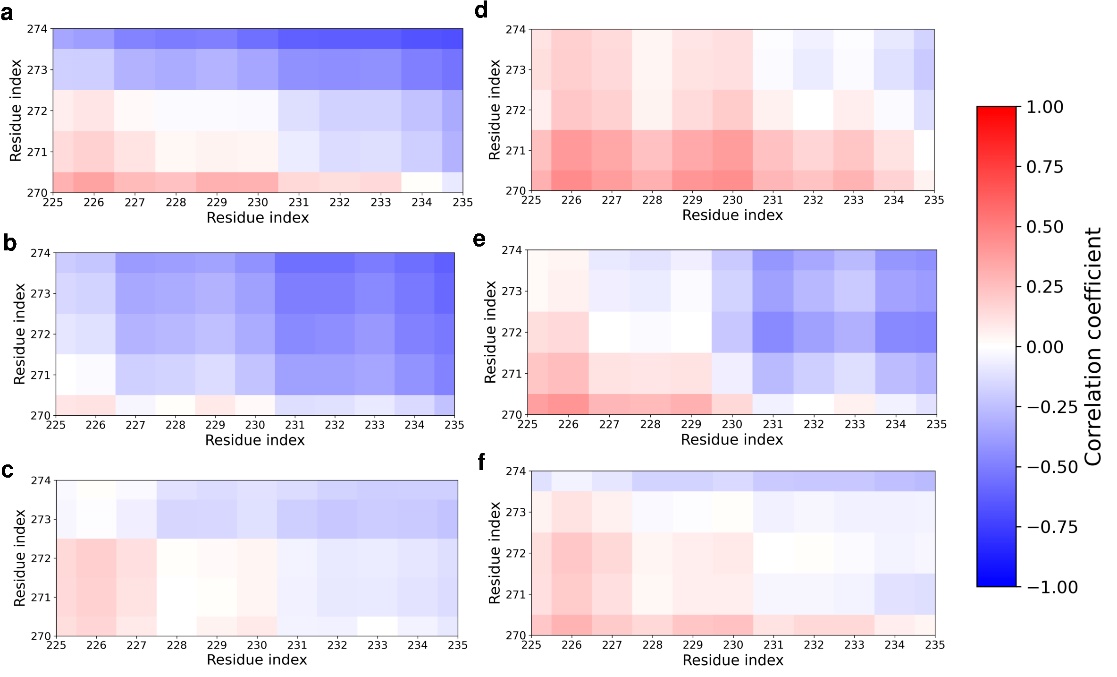

**Figure S18**: Dynamic Cross Correlation of BACE1 active site 2 (residue 225-235, including Asp228) and Insert D (residues 270-274) with (a) Atabecestat (b) Lanabecestat (c) Verubecestat (d) Bacopasaponin A (e) Bacopasaponin D (f) Bacopaside 1

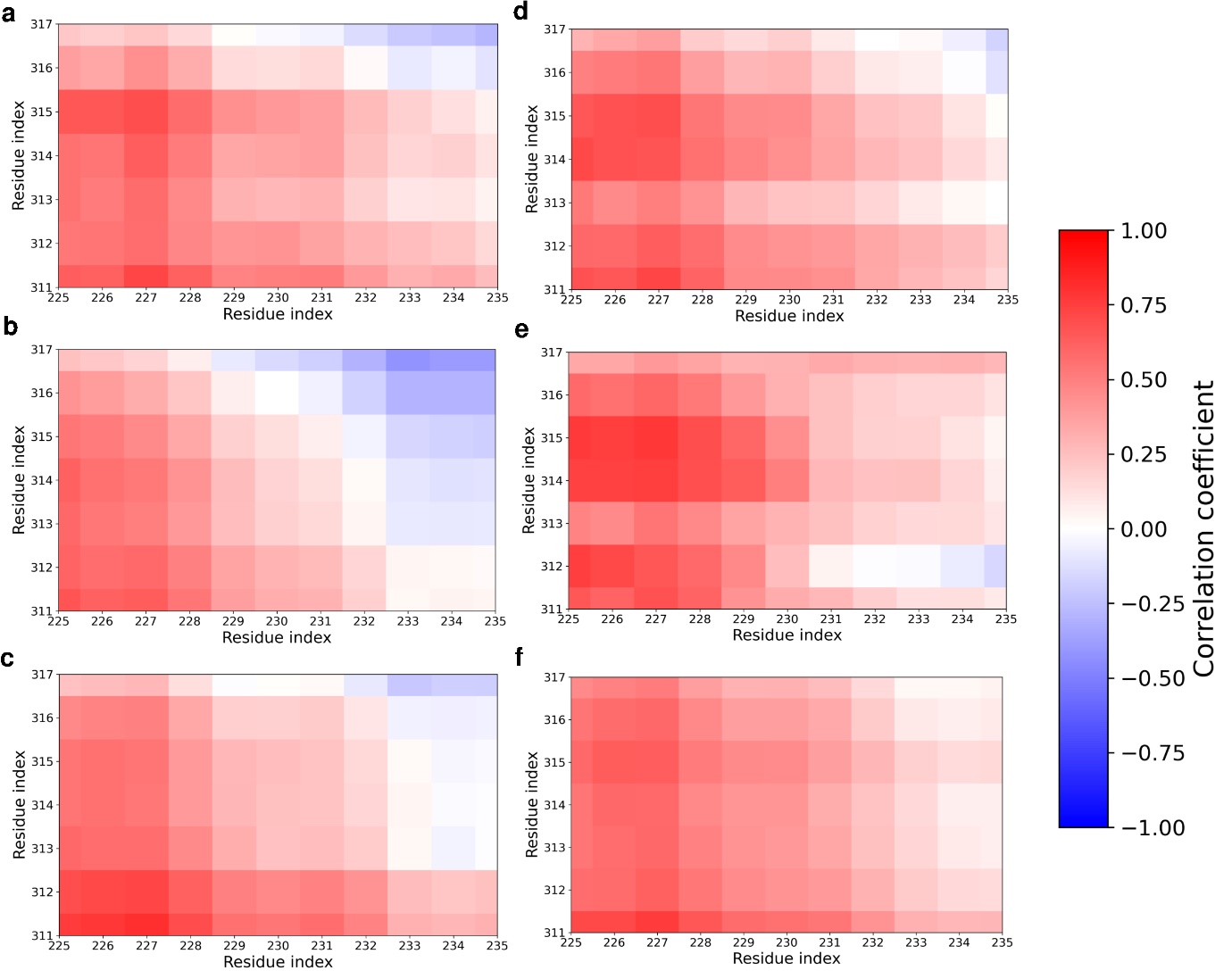

**Figure S19**: Dynamic Cross Correlation of BACE1 active site 2 (residue 225-235, including Asp228) and Insert F (residues 311-317) with (a) Atabecestat (b) Lanabecestat (c) Verubecestat (d) Bacopasaponin A (e) Bacopasaponin D (f) Bacopaside 1

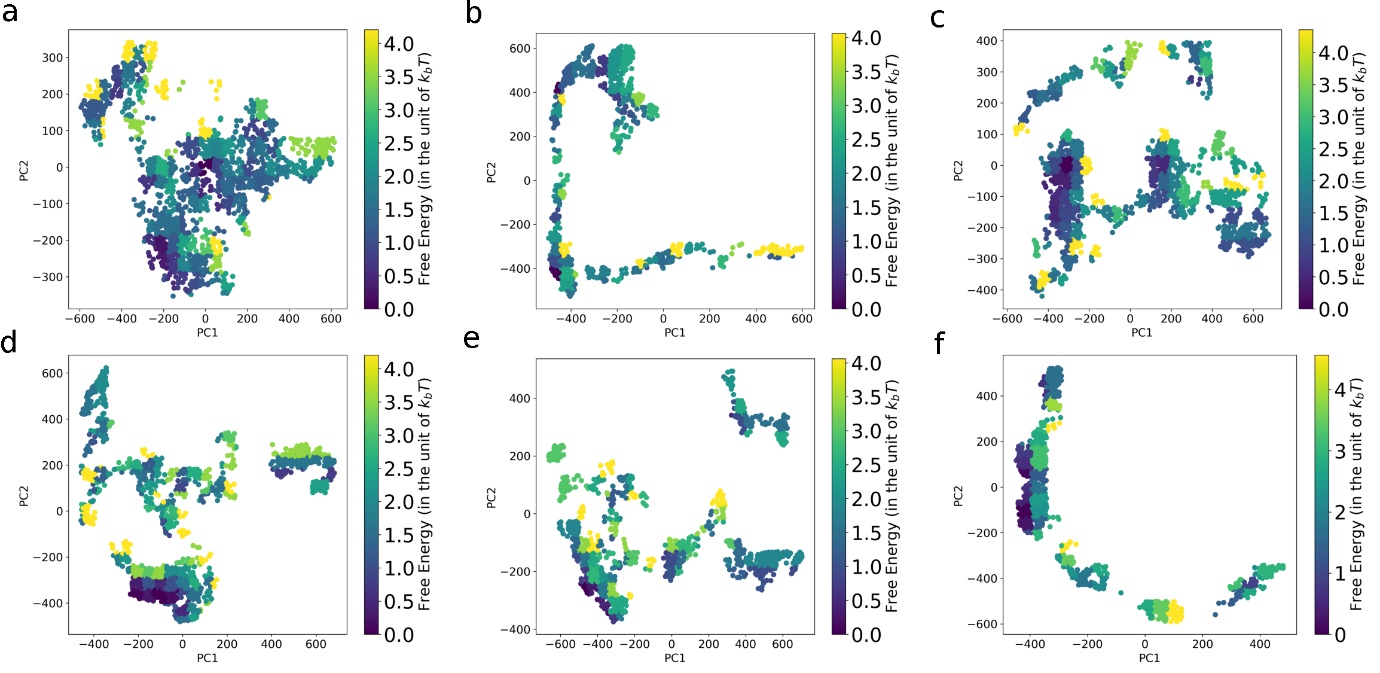

**Figure S20**: Free energy landscape in PCA space for BACE1 interaction with (a) Atabecestat (b) Lanabecestat (c) Verubecestat (d) Bacopasaponin A (e) Bacopasaponin D (f) Bacopaside 1

**Table S2**: Distance profile of Active site aspartates with important regions of BACE1 with respective ligand bindings

| Ligands | Asp32 | | | | | | Asp228 | | | | | |
| --- | --- | --- | --- | --- | --- | --- | --- | --- | --- | --- | --- | --- |
|  | R1 | R2 | R3 | R4 | R5 | R6 | R1 | R2 | R3 | R4 | R5 | R6 |
| Atabecestat | 13.50 | 15.57 | 15.48 | 33.20 | 37.96 | 36.62 | 14.69 | 14.99 | 23.11 | 33.89 | 34.69 | 35.79 |
| Lanabecestat | 15.77 | 16.27 | 15.68 | 23.16 | 38.71 | 28.85 | 15.04 | 15.78 | 19.59 | 22.45 | 35.22 | 27.46 |
| Verubecestat | 12.12 | 15.11 | 16.  29 | 21.73 | 37.123 | 36.41 | 12.99 | 13.06 | 20.85 | 22.44 | 32.87 | 34.  08 |
| Bacopasaponin A | 12.39 | 15.20 | 14.93 | 21.77 | 40.98 | 35.48 | 14.86 | 12.61 | 19.70 | 21.64 | 35.63 | 34.14 |
| Bacopasaponin D | 12.67 | 14.40 | 13.48 | 20.58 | 39.93 | 35.88 | 15.91 | 12.00 | 17.98 | 20.07 | 34.61 | 32.91 |
| Bacopaside 1 | 12.91 | 15.21 | 15.69 | 21.08 | 39.25 | 34.68 | 13.57 | 13.67 | 20.11 | 20.98 | 34.62 | 33.26 |
